# Deep learning enables quantitative subcellular analysis of plant-microbe interfaces

**DOI:** 10.64898/2026.02.05.703961

**Authors:** Stavros Korovesis, Shumei Wang, Lin Xu, Inès Giraudon, Daniela Rosales Hernandez, Enora Panek, Laure Boeglin, Maria-Myrto Kostareli, Max HJ Pluis, Baoyu Wang, Yuan Wang, Djenatte Abdennour, Harald Keller, Paul RJ Birch, Sebastian Schornack, Edouard Evangelisti

## Abstract

Specialized host-microbe interfaces are central to cellular interactions in plants. Intracellular structures such as haustoria formed by filamentous pathogens mediate nutrient exchange and effector delivery to host cells. Despite their biological importance, the lack of quantitative frameworks has largely confined the study of these interfaces to qualitative observations, limiting our ability to compare infection strategies, cellular responses, and spatial organization across cells and tissues. Here, we present [HFinder](https://github.com/EEvangelisti/hfinder), a deep learning-based framework for automated detection, segmentation, and quantitative analysis of plant-microbe interfaces in confocal images. Using an object-centric deep learning approach, HFinder enables robust identification of haustoria, microbial hyphae, and host organelles across diverse imaging conditions and pathosystems. We demonstrate that this framework supports quantitative analyses of subcellular processes at host-microbe interfaces, including effector secretion, perturbation of host cellular processes, and immune receptor accumulation at haustoria. HFinder provides a practical and scalable solution for the systematic digitalization of plant infection imaging data and establishes a general framework for quantitative studies of cellular dynamics at host-microbe contact zones.

## Introduction

Across plant-microbe interactions, specialized cell-cell interfaces create confined microenvironments for molecular exchange. In plants, such interfaces are prominently exemplified by specialized structures formed by filamentous microbes during intracellular colonization. For instance, mycorrhizal fungi establish hyphae-derived interfaces within plant cells, such as arbuscules and pelotons, to enable the exchange of mineral nutrients for photosynthetically derived carbon (Kuga *et al*., 2014; Zahn *et al*., 2023; Duan *et al*., 2024). Similarly, many fungal and oomycete plant pathogens produce hyphal protrusions into plant cells known as haustoria, which function as hubs for effector secretion and host manipulation (Szabo & Bushnell, 2001; Voegele *et al*., 2001; Hardham, 2007; Wang *et al*., 2018; Mapuranga *et al*., 2022). Despite their distinct evolutionary origins and biological outcomes, these interfaces share a common role as highly specialized platforms for molecular trafficking at the plant-microbe boundary.

Haustoria exhibit morphological diversity across taxa, but also within a given species, depending on host and environmental conditions. For instance, *Phytophthora* species typically form elongated, digitiform haustoria that may branch (Blackwell, 1953), whereas some downy mildews develop globose to lobate haustoria (Thines & Choi, 2016). Beyond microbial structures themselves, haustoria are associated with profound reorganization of host cell architecture. Several plant organelles, including nuclei, chloroplasts, and peroxisomes, reposition toward haustoria during infection (Lu *et al*., 2012; Wang *et al*., 2019; Savage *et al*., 2021), as do endomembrane compartments of the secretory pathway and exocyst subunits (Takemoto *et al*., 2003; Overdijk *et al*., 2020). This spatial rearrangement is paralleled by the focal accumulation of host immune receptors, such as the helper NLR NRC4 (Duggan *et al*., 2021), as well as multiple pathogen effector proteins (Bozkurt *et al*., 2011; Dagdas *et al*., 2018; Evangelisti *et al*., 2023). These interfaces therefore emerge as key sites of subcellular reorganization and immune signaling during infection.

Automated image analysis has become an essential component of modern cell biology, with deep learning approaches now widely adopted for microscopy data. Most frameworks for biological image analysis are built around dense, pixel-wise segmentation strategies, exemplified by U-Net and its derivatives, which predict a class label for every pixel in an image (Ronneberger *et al*., 2015). Such approaches have proven highly effective for segmenting continuous, homogeneous structures such as cells or tissues and have been extended to plant systems in both two- and three-dimensional contexts (Wolny *et al*., 2020; Greenwald *et al*., 2022). Instance-based methods such as Cellpose and StarDist build upon pixel-level predictions to reconstruct individual object instances (Schmidt *et al*., 2018; Stringer *et al*., 2021). Other architectures, including Mask R-CNN, combine object detection with instance segmentation and have been successfully applied to biological problems involving discrete objects (He *et al*., 2017). In parallel, a growing ecosystem of user-friendly pipelines and platforms has lowered the barrier to applying deep learning to microscopy, including Ilastik (Berg *et al*., 2019), ZeroCostDL4Mic (von Chamier *et al*., 2021), various cell and organelle segmentation pipelines (Hollandi *et al*., 2020; Aspert *et al*., 2022; Ding *et al*., 2025), and task-specific tools such as AMFinder and TAIM for quantifying mycorrhizal colonization (Evangelisti *et al*., 2021; Muta *et al*., 2022).

Most existing approaches are not tailored to the specific challenges posed by plant-microbe interfaces. Haustoria and associated subcellular features represent sparse, heterogeneous, anisotropic, and morphologically variable objects embedded within complex host tissues, often imaged across multiple fluorescence channels. To address these challenges, we developed HFinder, a deep learning-based framework designed for the analysis of plant-microbe interfaces in high-magnification confocal images. HFinder adopts an object-centric detection and segmentation strategy based on the YOLO architecture (Redmon *et al*., 2016), enabling efficient identification of discrete infection structures and organelles under realistic experimental conditions. Using successive training datasets, we demonstrate the versatility of the HFinder framework across distinct experimental scenarios, ranging from haustorium detection supported by biological reporters to datasets lacking such references, and to plant-pathogen systems exhibiting divergent haustorial morphologies. We further show that the framework can be readily extended to additional object classes, including plant organelles. Crucially, predictions are performed independently on individual imaging channels and subsequently reconciled through a decision scheme that incorporates user-defined biological constraints, reflecting the logic of multi-channel fluorescence microscopy. This channel-aware strategy enables quantitative analyses of molecular events at plant-microbe interfaces by explicitly encoding co-occurrence and mutual exclusivity relationships between subcellular signals. Together, HFinder provides a practical and scalable solution for the systematic digitalization and quantitative analysis of plant-microbe interaction images, addressing a critical gap at the interface of plant pathology, cell biology, and computational image analysis.

## Materials and methods

### Plant and microbial material

The *Nicotiana benthamiana* line used in this study is a laboratory cultivar originating from Australia (Bally *et al*., 2018). *Phytophthora palmivora* isolate LILI (accession no. P16830) was initially isolated from oil palm in Colombia (Torres *et al*., 2010) and maintained in the *Phytophthora* collection at Sophia Agrobiotech Institute (France). The transgenic *Phytophthora* lines used in this study, including LILI YFP-KDEL (Rey *et al*., 2015) and LILI Lifeact-mCitrine (Evangelisti *et al*., 2019), were maintained accordingly on medium supplemented with geneticin (G418) at 100 mg/L.

### Seed germination and growth conditions

*N. benthamiana* seeds were germinated on Neuhaus NPro substrate (Klasmann-Deilmann, Geeste, Germany) for 1 week at 24°C under a 16 h light / 8 h dark photoperiod. Seedlings were subsequently transferred to individual pots and grown for an additional 4 weeks under identical conditions. The biological control agents VectoBac WG (Valent BioSciences, Libertyville, IL, USA) and Entonem (Koppert, Rodenrijs, The Netherlands) were applied to the soil as recommended by the manufacturers. Transgenic *N. benthamiana* plants expressing the nuclear marker CFP-NbH2B (histone H2B fused to cyan fluorescent protein) were grown under controlled glasshouse conditions using the same photoperiod.

### Confocal imaging

Images used to train HFinder were acquired using either a Zeiss LSM 880 laser-scanning confocal microscope (Zeiss, Oberkochen, Germany) equipped with 20× dry and 63× water-immersion objectives and an argon laser, or a Leica TCS SP8 laser-scanning confocal microscope (Leica, Wetzlar, Germany) equipped with 25× and 63× water-immersion objectives and a white-light laser (470-670 nm) compatible with a broad range of fluorophores. Wavelengths of 470, 488, 514, and 581 nm were used for excitation of mTFP1, eGFP, mCitrine, and red fluorescent proteins (tdTomato, DsRed, or oScarlet), respectively. The oScarlet reporter used in this study is a derivative of mScarlet-I carrying the E95D mutation, which reduces mistargeting to lysosomes (Fenno *et al*., 2020). For effector translocation experiments, images were obtained using a Nikon A1plus laser-scanning confocal microscope (Nikon, Tokyo, Japan) equipped with a 40× water-immersion objective. CFP fluorescence was excited at 456 nm and collected between 464-500 nm, while mRFP fluorescence was excited at 561 nm and collected between 570-620 nm. The pinhole was set to 1.2 Airy units to optimize optical sectioning.

### Software design

#### Implementation

HFinder provides an automated pipeline for the detection and delineation of haustoria and hyphae, as well as plant nuclei and chloroplasts, in multi-channel confocal images. It is implemented as a command-line tool with dedicated modules for dataset preprocessing (`hfinder preprocess`), training of a lightweight YOLOv8 convolutional neural network designed for simultaneous instance detection and delineation (YOLOv8n-seg; `hfinder train`), and YOLO-derived instance and semantic segmentation (`hfinder predict`). The software is written in Python building on the [PyTorch](https://pytorch.org/) framework through the [Ultralytics](https://github.com/ultralytics/ultralytics) [YOLO](https://github.com/ultralytics/ultralytics) implementation. HFinder runs on Microsoft Windows, macOS, and GNU/Linux.

#### Analysis pipeline

A typical HFinder workflow begins with the assembly of a training dataset consisting of a directory of TIFF images accompanied by a JSON configuration file specifying how meaningful signals should be extracted. The next step involves dataset preprocessing, which enables manual inspection of the contours generated by HFinder and fine-tuning of the signal extraction strategy. Once a suitable dataset has been established, model training is performed, producing a trained model together with performance metrics that allow users to assess training quality. The trained model can then be used for predictions, which can be further analysed with auxiliary scripts provided for prediction visualization (`annot2images.py`), signal extraction (`annot2signal.py`), and distance measurements (`annot2distances.py`). For convenience, HFinder is distributed with pre-trained models that can be applied directly to confocal image analysis (available on Zenodo: https://doi.org/10.5281/zenodo.17091805). However, given the diversity of confocal imaging conditions, biological structures, magnifications, and subcellular signals, these pre-trained models may require fine-tuning before application to other experimental systems.

### Deep learning

#### Training dataset and classes

We assembled a training dataset of 580 single-plane and Z-stack confocal TIFF images, publicly available on Zenodo **(Table 1, Supporting Table 1)**. Each image contained between 2 and 5 channels, capturing signals from fluorescent proteins targeted either to a single compartment (e.g., nucleus) or to multiple compartments (e.g., nucleus and plasma membrane). In most cases, chlorophyll autofluorescence was included to visualize chloroplasts. For images derived from infected plant material, fluorescently labelled *Phytophthora palmivora* strains were used to reveal the hyphal network.

**Table 1.**
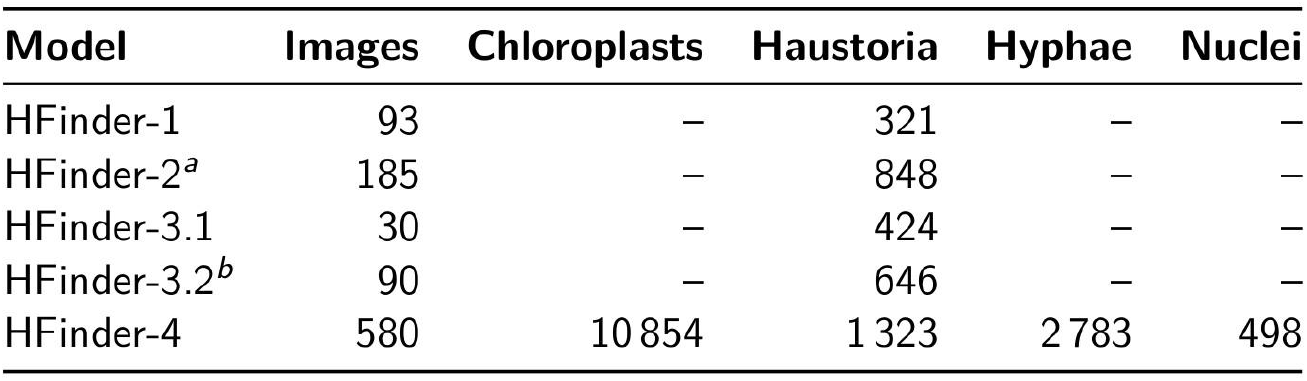
Summary of training images and annotated instances for HFinder models. Each model was trained on a distinct image dataset. The table reports the number of training images and the corresponding number of annotated biological structures used during training. ^*a*^: the HFinder-2 training dataset includes all images from the HFinder-1 dataset, supplemented with 92 additional images corresponding to 527 extra haustoria annotated by transfer learning. ^*b*^: The HFinder-3.2 training dataset extends the HFinder-3.1 dataset by adding 60 images from the HFinder-2 dataset, corresponding to 222 haustoria from *Phytophthora* infections.

#### Image preprocessing

Per-channel thresholding used scikit-image methods (“isodata,” “li,” “otsu,” “yen,” “triangle”) or a numeric threshold; an “auto” mode selected Otsu or Triangle based on image skewness. Binary masks obtained by thresholding, semi-automated annotation, or manual curation using Makesense.ai (Skalski, 2019) were then converted into polygonal annotations compatible with YOLO segmentation, represented as flattened lists of normalized vertex coordinates [x_1_, y_1_, …, x_*n*_, y_*n*_] in the [0, 1] range. Following initial vectorization, contours with an area below a user-defined threshold (`min_area`), computed in pixel space prior to normalization, were discarded to remove small spurious objects and noise. The remaining polygonal contours were then subjected to a standardized geometric post-processing pipeline to reduce contour complexity and to homogenize vertex density across instances. Each polygon was first simplified using the Douglas-Peucker algorithm implemented in OpenCV (`cv2.approxPolyDP`). The simplification tolerance *ε* was defined relative to the contour perimeter using a user-defined parameter (`epsilon_rel`, default: 0.001), such as *ε* = `epsilon_rel` × *P*, where *P* is the polygon perimeter computed with `cv2.arcLength`. To prevent excessive simplification, polygons yielding fewer than a minimum number of vertices (`min_points`) after approximation were discarded in favor of the original, unsimplified contour. Simplified polygons were subsequently resampled to generate vertices distributed at approximately uniform arc-length intervals along the contour. Polygons were explicitly closed, cumulative arc lengths were computed along successive segments, and new vertices were interpolated linearly at fixed distances. The target number of vertices was set to `target_points` (default: 120), with an effective upper bound *K* = `min(target_points, max(`*n*`, 4))` (where *n* denotes the number of vertices of the polygon; the lower bound of 4 ensures a minimally valid closed polygon) to avoid over-representation of small instances. This resampling step ensures comparable vertex densities across objects of different sizes and shapes while preserving overall contour geometry. All parameters controlling polygon simplification and resampling (`epsilon_rel`, `min_points`, `min_area`, and `target_points`) were defined globally via the HFinder configuration to ensure consistency across datasets and preprocessing runs.

#### Classifier parameters and training

Four versions of HFinder were generated in this study. The initial model, HFinder-1, was trained from naïve YOLOv8 nano segmentation weights (`yolov8n-seg`) using default training parameters (200 epochs, image size 640 × 640 pixels, and a 20% validation split). HFinder-2 was trained using the same settings, with weights initialized from HFinder-1. HFinder-3.1 and HFinder-3.2 were likewise trained for 200 epochs, using the same base architecture and training parameters, but with modified preprocessing settings (`min_points` = 3 and `min_area` = 5) to accommodate morphology-specific fine-tuning. HFinder-4 was trained from naive YOLOv8 nano segmentation weights using 300 epochs and otherwise default training parameters.

#### Classifier evaluation

Model training was monitored using the standard YOLOv8 segmentation objective, which comprises four loss components: bounding box regression loss, segmentation mask loss, classification loss, and distribution focal loss (Redmon *et al*., 2016; Jocher *et al*., 2023). Convergence of these loss terms during training was used to assess optimization stability and to detect potential overfitting.

In parallel, model performance on the validation subset was quantified using precision, recall, and mean Average Precision (mAP). Precision and recall were used to evaluate detection specificity and sensitivity, respectively. Precision was defined as the proportion of true positive detections among all predicted positives, and recall as the proportion of true positive detections among all ground-truth objects:

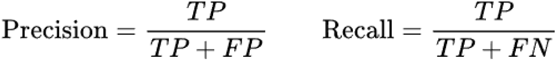

where *TP* denotes true positives, *FP* false positives, and *FN* false negatives.

Detection and segmentation performance were further summarized using mean Average Precision (mAP), which integrates precision-recall curves over confidence thresholds. Average Precision (AP) was computed for each class at a given intersection-over-union (IoU) threshold, and mAP was obtained by averaging AP values across classes. mAP50 corresponds to AP computed at a fixed IoU threshold of 0.5, whereas mAP50-95 represents the mean AP averaged over IoU thresholds ranging from 0.5 to 0.95 in increments of 0.05, providing a more stringent and comprehensive measure of model performance (Redmon *et al*., 2016; Jocher *et al*., 2023).

For independent evaluation on held-out test datasets, model performance was assessed using precision and recall, together with the IoU metric computed between predicted and expert-annotated instance masks. IoU was defined as:

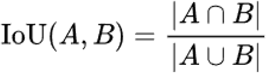

where *A* and *B* denote the predicted and reference instance masks, respectively.

## Results

### HFinder enables robust detection and instance segmentation of *Phytophthora* haustoria

Haustoria formed by *Phytophthora* species during plant infection act as secretion hubs for effectors and other virulence-associated proteins (Wang *et al*., 2018). Because these structures frequently resemble emerging hyphae, their manual annotation is both challenging and error-prone (Guyon *et al*., 2025), particularly in low-contrast imaging conditions. To establish a high-confidence haustorium training reference, we first curated a set of biotrophy images leveraging NRC4 signal enrichment at haustorial sites (Duggan *et al*., 2021) **(Fig. S1, Table 1, Supporting Table 1)**. We then trained an initial model, hereafter termed HFinder-1, to detect haustoria from raw multi-channel confocal images **(Fig. 1, S2, S3)**.

**Figure 1.**
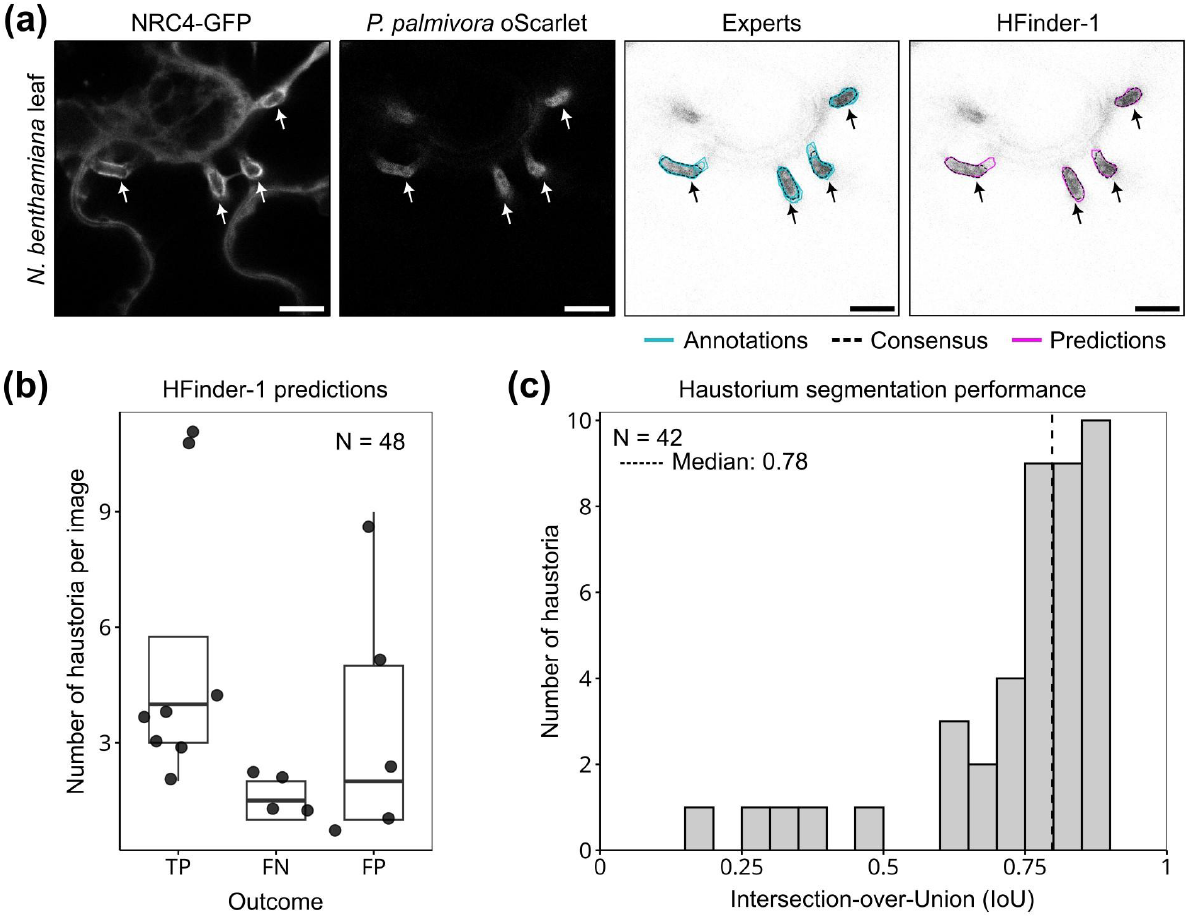
HFinder-1 enables robust quantification and segmentation of *Phytophthora* haustoria. **(a)** Representative confocal images of NRC4-expressing *Nicotiana benthamiana* leaves inoculated with a transgenic *Phytophthora palmivora* strain expressing a cytoplasmic oScarlet. Expert-annotated haustoria (cyan) were used to derive a consensus contour (black dashed line), against which HFinder-1 predictions (magenta) were compared. Bounding boxes are omitted for clarity. Arrows indicate haustoria. Scale bars: 10 µm. **(b)** Distribution of HFinder-1 detection outcomes as true positives (TP), false negatives (FN), and false positives (FP). Each point represents the number of haustoria per image (48 haustoria across nine images). **(c)** Intersection-over-Union (IoU) values for the 42 true-positive haustoria segmented by HFinder-1. The median IoU was 0.78 (dashed line).

Channels not used for annotation were provided as additional background information. The training dataset comprised 321 manually annotated haustoria spanning a wide spatial distribution across fields of view **(Fig. S1b, Supporting Table 1)** and exhibiting a relatively narrow size range consistent with canonical *Phytophthora* haustorial morphology, thereby limiting morphological ambiguity during training **(Fig. S1c)**. Upon training, model optimization converged rapidly, as indicated by steadily decreasing bounding-box regression, segmentation-mask, classification, and distribution focal losses, without evidence of overfitting **(Fig. S2a)**. Concomitantly, HFinder-1 displayed progressive gains in precision, recall, and mAP50 on the validation set, demonstrating effective generalization **(Fig. S2b)**. To independently assess model performance, we assembled a test set of 48 NRC4-positive haustoria extracted from 9 independent confocal images deliberately selected for their weak oScarlet reporter signal, thereby imposing fuzzy object boundaries and mimicking suboptimal imaging conditions **(Fig. 1a)**. HFinder-1 correctly identified 42 haustoria (87.5%; mean confidence score: 0.70), missed 6 cases (false negatives), and produced 18 additional predictions lacking corresponding NRC4 enrichment (false positives; mean confidence score: 0.50) **(Fig. 1b)**. Because HFinder produces instance-level segmentations, performance was additionally assessed using the distribution of intersection-over-union (IoU) values between predicted and expert-annotated haustoria. The median IoU was 0.78, with only five predictions falling below 0.50 **(Fig. 1c)**. These lower-scoring cases reflected partial rather than incorrect segmentations and frequently coincided with substantial inter-expert variability **(Fig. S3)**. Together, these results demonstrate that deep learning-based approaches can robustly identify and segment oomycete haustoria in high-magnification confocal microscopy images.

### Transfer learning improves haustorium detection confidence and spatial focus

We next asked whether transfer learning could improve not only detection sensitivity, but also the quality of haustorium recognition in terms of confidence calibration and spatial specificity. To this end, we extended the original HFinder-1 training dataset with images acquired under additional experimental conditions, including different microscopes, fluorescent markers, and independent laboratories **(Fig. 2, S4-S6)**. These images were initially annotated using HFinder-1 predictions, followed by manual removal of obvious misannotations. The resulting curated dataset comprised 92 additional images featuring 527 new haustoria **(Supporting Table 1)** and was used to fine-tune the model, yielding HFinder-2 initialized from HFinder-1 weights **(Fig. S4)**. Training of HFinder-2 showed stable convergence of all four components of the YOLOv8 objective function **(Fig. S4a)**, and the model reached maximal fitness after 77 epochs **(Fig. S4b)**. We then compared HFinder-1 and HFinder-2 on an expanded test set combining the original validation images with nine additional images drawn from the transfer-learning dataset. Relative to HFinder-1, HFinder-2 exhibited a clear reduction in false negatives **(Fig. S5a**,**c)**, resulting in a significant increase in recall **(Fig. 2b)**. Precision showed a consistent upward trend associated with a reduction in false positives **(Fig. S5a**,**c)**, although this increase did not reach statistical significance **(Fig. 2b)**.

**Figure 2.**
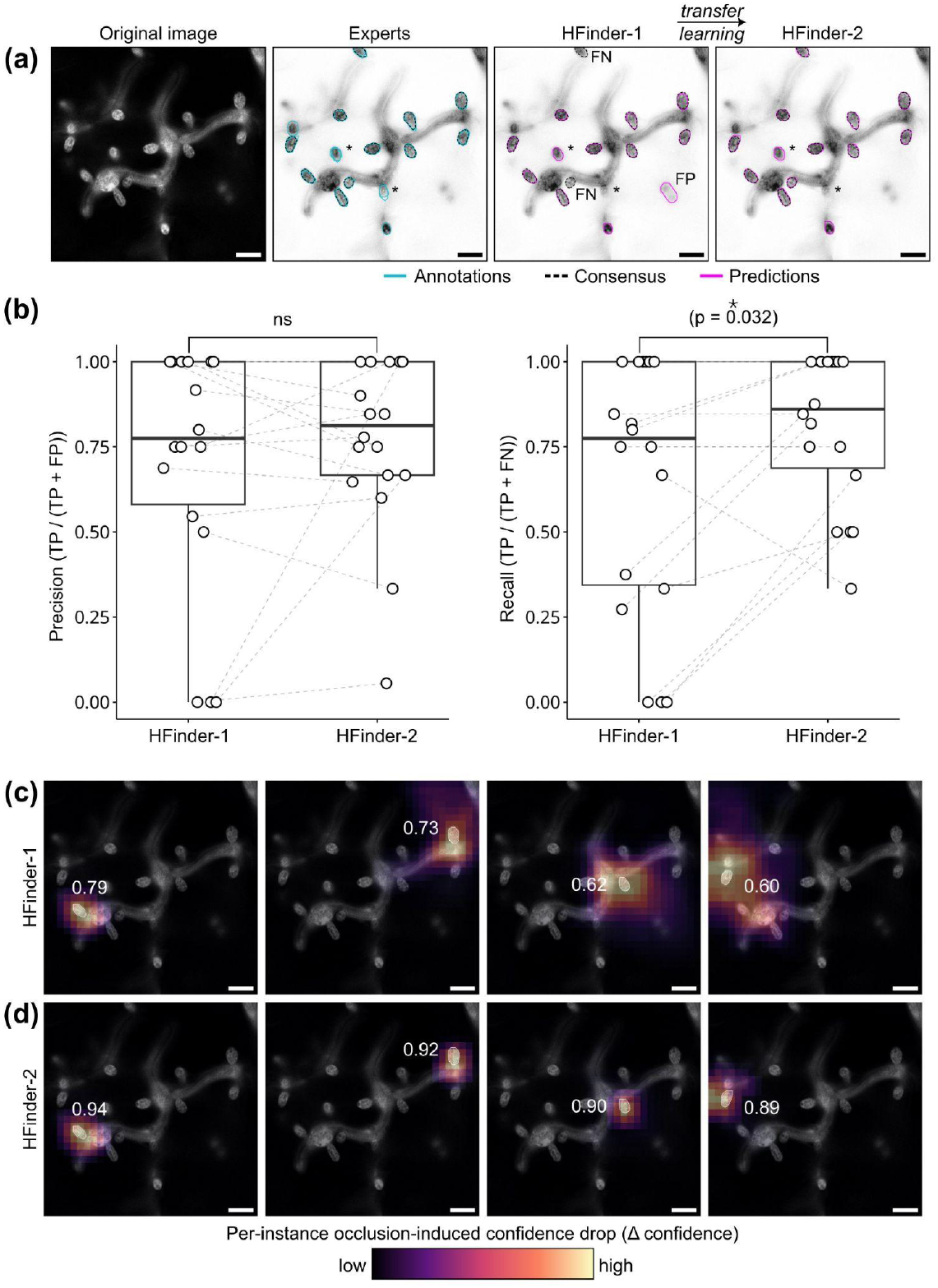
Transfer learning improves haustorium detection confidence and spatial focus. **(a)** Representative example illustrating the effect of transfer learning from HFinder-1 to HFinder-2 on *Phytophthora palmivora* haustoria detection. Expert annotations (cyan) and consensus contours (black dashed lines) are compared with predictions from HFinder-1 and HFinder-2 (magenta). Asterisks (*) indicate ambiguous cases excluded from quantitative analyses. **(b)** Precision (left) and recall (right) of HFinder-1 and HFinder-2 measured on the haustoria test image set. Each point corresponds to one image. Grey dashed lines indicate pairs. Statistical significance was assessed using paired Wilcoxon signed-rank tests; p-values are indicated. **(c-d)** Occlusion sensitivity analysis for HFinder-1 **(c)** and HFinder-2 **(d)**. Square occlusions were systematically applied across the input image, and the resulting decrease in detection confidence was quantified for each tracked haustorium instance. Heatmaps represent the spatial distribution of the occlusion-induced confidence decrease (Δ confidence) for four representative haustoria. Numbers in white indicate the baseline detection confidence measured on the unoccluded image. Scale bars: 10 µm. ns: not significant; TP: true positive; FP: false positive; FN: false negative.

While improvements in global performance metrics were modest, qualitative differences between the two models were evident. Occlusion sensitivity analysis revealed a pronounced reduction in the spatial extent of image regions contributing to haustorium detection in HFinder-2 compared with HFinder-1 **(Fig. 2c,d)**. In HFinder-1, confidence decreases were distributed over broader image regions, often encompassing surrounding hyphae or background signal **(Fig. 2c)**. In contrast, HFinder-2 displayed sharply localized confidence drops centered on haustorial structures, indicating a shift toward more spatially focused and biologically meaningful feature utilization **(Fig. 2d)**. Consistent with this improved spatial focusing, HFinder-2 produced significantly higher confidence scores for true positive detections compared with HFinder-1 (paired Wilcoxon signed-rank test, p = 0.014), whereas confidence scores associated with false positive predictions remained unchanged (p = 0.81) **(Fig. S6)**. Together, these results demonstrate that transfer learning primarily enhances the calibration and spatial specificity of haustorium recognition, enabling HFinder to evaluate and localize haustoria more confidently and interpretably without compromising generalization.

### HFinder detects and segments haustoria across diverse plant-oomycete pathosystems

Whereas haustoria formed by *Phytophthora* species are typically digitiform, other oomycetes produce haustoria with markedly distinct morphologies. In particular, members of the genus *Hyaloperonospora* develop lobate to globose haustoria (Thines & Choi, 2016). We therefore asked whether HFinder could generalize to such divergent morphologies and whether targeted fine-tuning could further improve detection performance. To maintain imaging conditions comparable to those used previously, we acquired confocal images of *Arabidopsis thaliana* leaves infected with *Hyaloperonospora arabidopsidis* and stained with Trypan blue, exploiting the intrinsic fluorescence of the dye (Hoffmeister *et al*., 2020). We then compared the performance of HFinder-2 with two fine-tuned variants: HFinder-3.1, retrained exclusively on an *H. arabidopsidis* haustoria dataset (N = 424 haustoria), and HFinder-3.2, retrained on a combined dataset containing both *H. arabidopsidis* and *P. palmivora* haustoria (N = 646 haustoria, including 222 haustoria from *Phytophthora*) **(Fig. 3, Table 1, Supporting Table 1)**. Only images for which predictions were produced by all compared models were retained for per-image precision and recall analyses.

**Figure 3.**
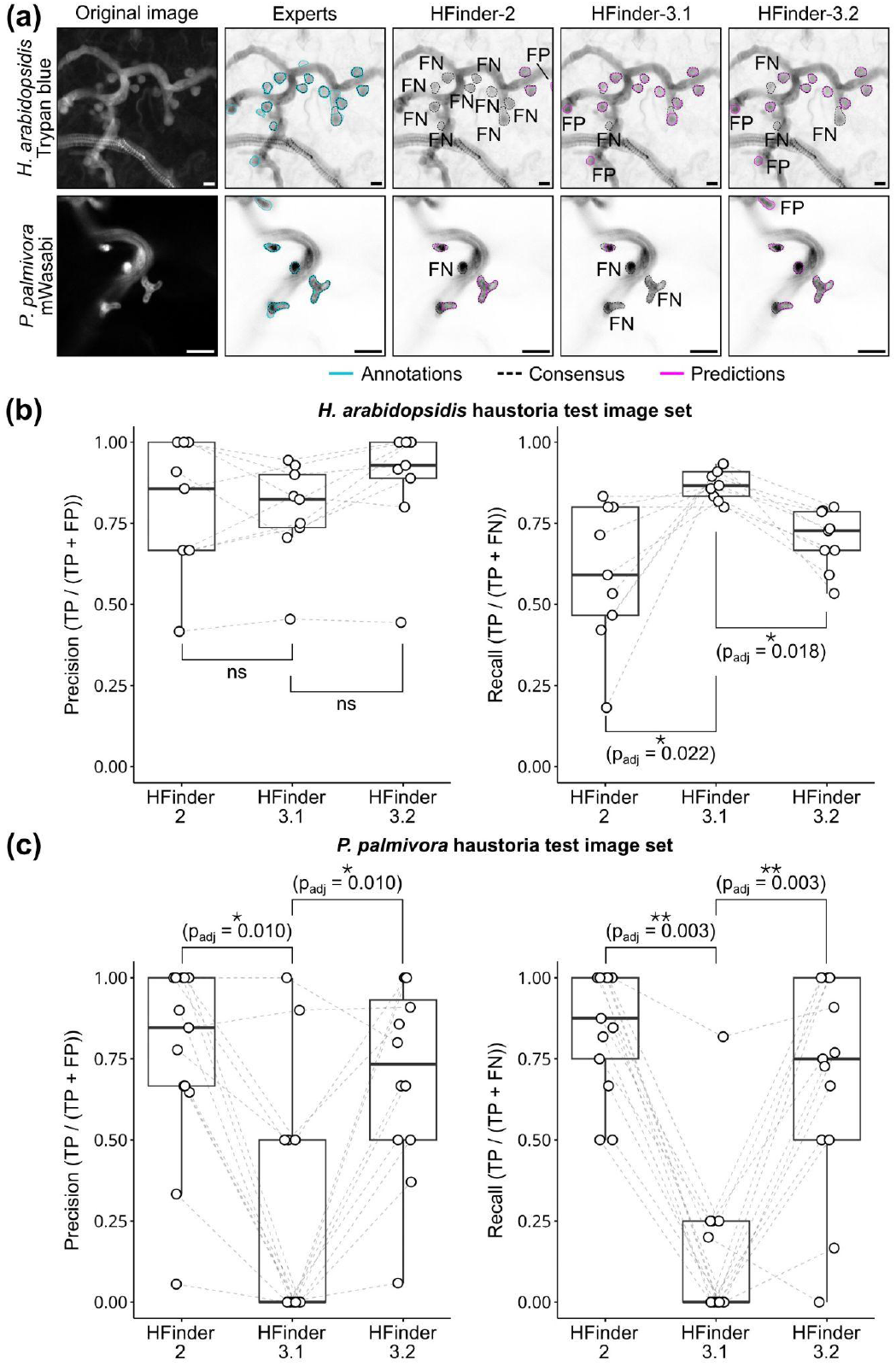
Morphology-specific fine-tuning reveals trade-offs between haustorium detection performance and generalization. **(a)** Representative confocal images illustrating haustorium detection across successive HFinder versions. Infected leaves of *Arabidopsis thaliana* inoculated with *Hyaloperonospora arabidopsidis* (top row, trypan blue staining) and *Nicotiana benthamiana* infected with *Phytophthora palmivora* (bottom row, mWasabi-labelled strain) are shown. Expert annotations (cyan) and inter-expert consensus contours (black dashed lines) are compared with predictions generated by HFinder-2, HFinder-3.1, and HFinder-3.2 (magenta). False positives (FP) and false negatives (FN) are indicated. Scale bars: 10 µm. **(b-c)** Precision (left) and recall (right) measured for HFinder-2, HFinder-3.1, and HFinder-3.2 on the *H. arabidopsidis* **(b)** and *P. palmivora* **(c)** haustoria test image sets. Each point corresponds to one test image, with paired measurements connected by thin lines. Planned paired comparisons were assessed using Wilcoxon signed-rank tests with Holm correction; adjusted p-values are indicated.

On the *H. arabidopsidis* test set, HFinder-2 achieved a relatively high mean precision (0.80) but produced a substantial number of false negatives, resulting in a modest mean recall of 0.59. This behavior is consistent with the more globular morphology of *H. arabidopsidis* haustoria, which contrasts with the predominantly digitiform shapes encountered during the initial *Phytophthora*-focused training **(Fig. 3a, b)**. Fine-tuning on *H. arabidopsidis* haustoria (HFinder-3.1) did not significantly affect precision (mean precision: 0.79), but led to a significant improvement in recall, which increased to a mean of 0.87 **(Fig. 3b)**. The compromise model HFinder-3.2 exhibited comparable precision (mean: 0.88) and an intermediate recall (mean: 0.70), consistent with its mixed training strategy. We next evaluated all three models on the *P. palmivora* test set **(Fig. 3c)**. As previously observed, HFinder-2 performed well on this dataset, with high mean precision (0.74) and recall (0.84). In contrast, the *H. arabidopsidis*-specialized model HFinder-3.1 showed a pronounced loss of performance, with mean precision and recall dropping to 0.24 and 0.14, respectively. The compromise model HFinder-3.2 displayed intermediate performance (mean precision: 0.69; mean recall: 0.69), which did not significantly differ from that of HFinder-2. Together, these results demonstrate that fine-tuning HFinder on morphology-specific datasets substantially improves haustorium detection within the corresponding pathosystem, primarily by reducing false negatives and increasing recall. However, these gains are accompanied by a loss of performance on previously learned morphologies. This trade-off can be mitigated by using mixed training datasets, which preserve generalization while incurring only a modest reduction in detection performance.

### Multi-class HFinder extends detection to hyphal networks and plant organelles

To extend HFinder toward a more comprehensive analysis of infected plant tissues, we developed HFinder-4, a multi-class model trained to simultaneously detect haustoria, hyphae, and plant organelles reported to relocalize to haustoria, including nuclei (Caillaud *et al*., 2012; Sharma & Chandran, 2022) and chloroplasts (Savage *et al*., 2021). The training dataset comprised 580 multi-channel confocal images with semi-supervised annotations for 10,854 chloroplasts, 1,323 haustoria, 2,783 hyphae, and 498 nuclei **(Fig. S7, Supporting Table 1)**. Annotation of this dataset combined manual curation with semi-automatic mask generation based on user-defined thresholding, providing a pragmatic compromise between annotation accuracy and scalability. These segmentation masks were subsequently used to derive the ground-truth annotations. To ensure balanced learning across classes, HFinder-4 was trained using identical class proportions in the training and validation sets. Nevertheless, strong instance-level imbalance (intrinsic to the biological abundance of the objects of interest) was deliberately preserved, allowing model performance to be assessed under conditions closely reflecting real experimental datasets. During training, HFinder-4 exhibited a steady decrease in loss accompanied by a progressive increase in evaluation metrics, reaching optimal performance after approximately 240 training epochs **(Fig. S8)**.

Model performance was next evaluated on an independent test dataset comprising images of nuclei (labelled with fluorophores carrying a nuclear localization signal), chloroplasts (detected via chlorophyll autofluorescence), and a transgenic *P. palmivora* strain expressing a cytoplasmic fluorescent marker labelling hyphae and haustoria. For plant nuclei, both precision and recall approached unity (> 0.95), reflecting robust segmentation of these globular, homogeneous structures under our imaging conditions **(Fig. 4a)**. Chloroplast segmentation achieved precision and recall values close to 0.7, indicative of good performance despite the semi-supervised nature of the training annotations **(Fig. 4b)**. Frequent merging of adjacent chloroplasts was observed when instances were closely spaced **(Figs. 4b, S9)**, a consequence of training masks that often grouped neighboring chloroplasts due to the absence of exhaustive manual instance separation. Detection of haustoria and hyphae was characterized by recall values close to 1, indicating that HFinder-4 rarely failed to detect these structures **(Fig. 4c)**. In contrast, mean precision was 0.60 for haustoria and below 0.50 for hyphae. This reduction in precision correlated with discrepancies between expert annotations used to generate consensus masks, reflecting the intrinsic difficulty of defining hyphal boundaries in single optical sections. In particular, signal attenuation caused by hyphae moving out of the focal plane when analyzing single optical sections rather than maximum-intensity projections introduces ambiguity in instance delineation.

**Figure 4.**
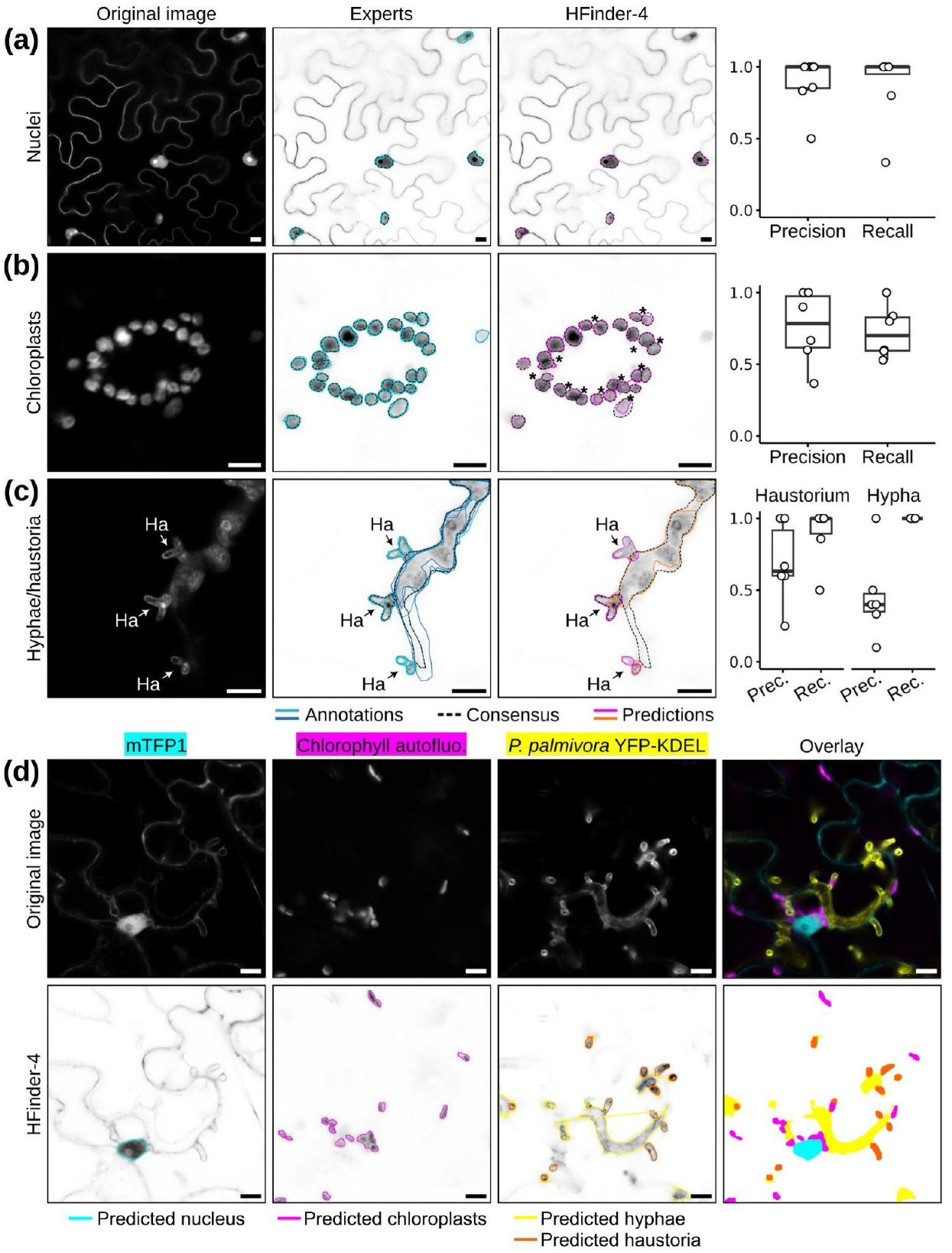
HFinder simultaneously labels and segments infection structures and plant organelles. **(a-c)** Representative confocal images showing HFinder-4 segmentation of plant nuclei **(a)**, chloroplasts **(b)**, and oomycete hyphae with haustoria **(c)**. Original images (left), expert annotations and consensus (middle), and HFinder-4 predictions (right) are shown for each class. Right-hand panels display segmentation performance, quantified as precision and recall based on intersection-over-union (IoU) between expert consensus annotations and HFinder predictions. Each dot represents one independently annotated image. Stars indicate merged chloroplast instances predicted as single objects. **(d)** Representative multi-channel confocal image used to evaluate channel attribution by HFinder-4, showing nuclei (mTFP1), chlorophyll autofluorescence, and *Phytophthora palmivora* YFP-KDEL signals, together with the corresponding HFinder predictions and overlay. Scale bars: 10 µm. Ha, haustorium; autofluo., autofluorescence; Prec., precision; Rec., recall.

Beyond the increased number of object classes, HFinder-4 introduces the additional challenge of correctly assigning predictions to the appropriate imaging channels. We found that channel attribution was performed accurately and consistently **(Fig. 4d, S10)** through a post-processing strategy that combines YOLO prediction confidence scores with a rule-based assignment framework. This framework explicitly accounts for co-occurring classes (e.g. hyphae and haustoria) and mutually exclusive classes (e.g. chloroplasts and hyphae), enabling reliable multi-channel interpretation **(Fig. S10)**. Together, these results demonstrate that HFinder-4 can simultaneously segment plant organelles and microbial infection structures across multiple imaging channels, providing a robust foundation for the systematic digitalization and quantitative analysis of plant-pathogen interaction images.

### HFinder enables quantitative analysis of molecular events at subcellular resolution in infected plant cells

We next asked whether HFinder could be leveraged to quantify plant-microbe interfaces at the subcellular scale **(Fig. 5)**. As a proof of principle, we quantified the enrichment of the helper NLR NRC4 at the haustorial periphery during *P. palmivora* infection of *N. benthamiana*. We reused HFinder-2 predictions obtained on a test dataset of NRC4-GFP-expressing leaves, in which 69 haustoria were automatically detected and segmented. The predicted haustorial outlines were used as masks to extract local GFP intensities reporting NRC4 distribution **(Fig. 5a)**. For each haustorium, we calculated the ratio between the mean GFP signal within the haustorial region and a local baseline signal, obtained by automatic exclusion of haustorial areas from the GFP channel. On average, NRC4-GFP intensity at haustoria was 7.6-fold higher than background levels, with half of the detected haustoria falling within the 93rd percentile of the global GFP intensity distribution **(Fig. 5a)**. These results demonstrate a robust and reproducible enrichment of NRC4 at the haustorial periphery, consistent with its preferential accumulation at this interface.

**Figure 5.**
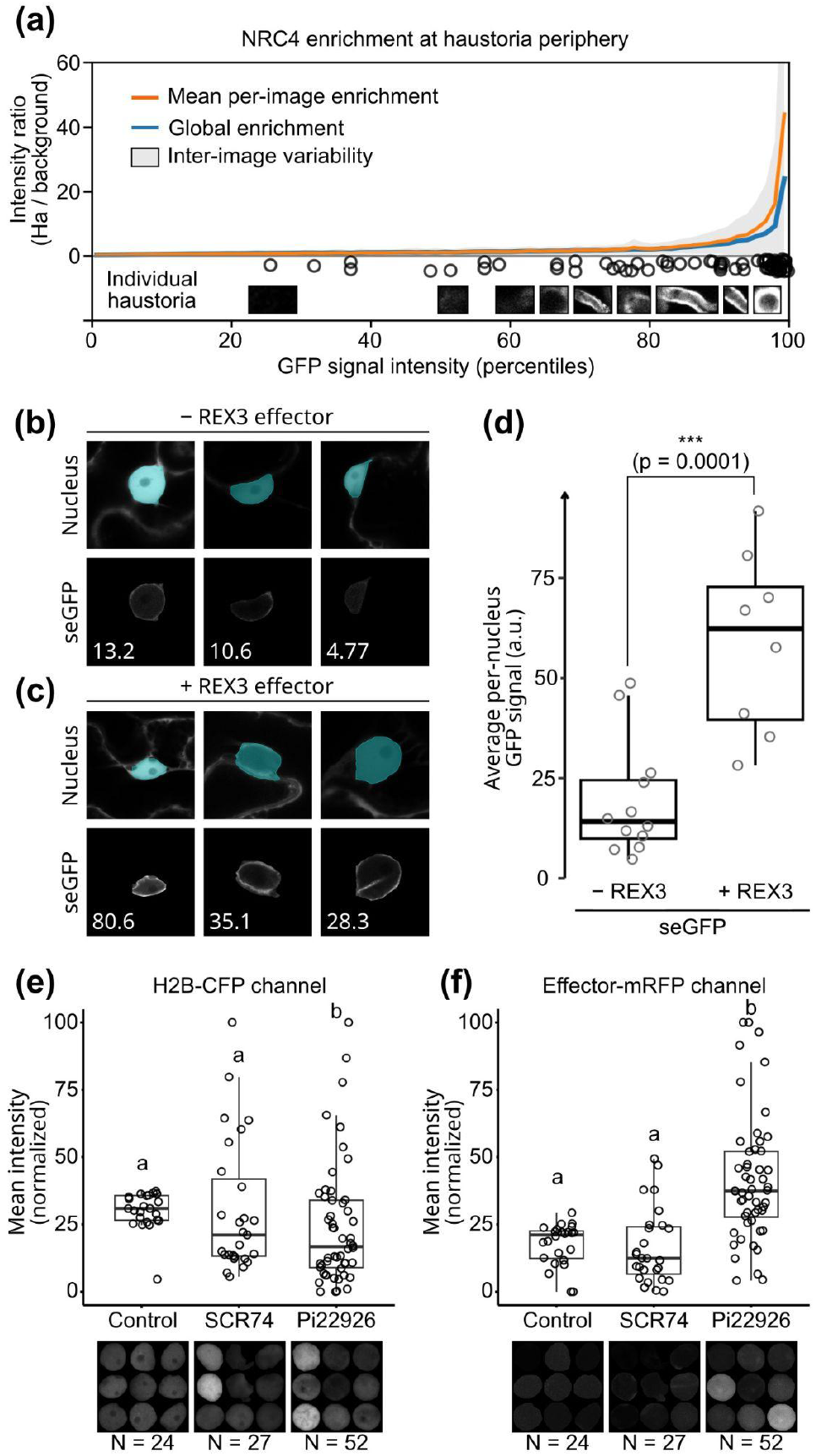
HFinder enables quantitative analysis of molecular events at the subcellular level. **(a)** Enrichment of NRC4-GFP at the haustorial periphery, quantified as the ratio of fluorescence intensity measured within haustorial regions relative to the surrounding background. Enrichment values are plotted as a function of GFP signal intensity percentiles. Black circles represent the mean enrichment value of individual haustoria. The orange curve corresponds to the mean per-image enrichment, the blue curve to the global enrichment across all images, and the grey shaded area indicates inter-image variability. Insets show representative examples of haustorial regions automatically detected and extracted by HFinder. **(b-c)** Representative confocal images illustrating plant cell nuclei automatically segmented by HFinder (cyan) in *Nicotiana benthamiana* cells co-expressing a nuclear-cytoplasmic DsRed marker and a GFP fused to the Pathogenesis-Related Protein 1 signal peptide (seGFP), expressed either alone (−REX3; **b**) or together with the RXLR effector REX3 (+REX3; **c**). Lower panels show the corresponding automatically extracted perinuclear seGFP signal for each nucleus, with the average fluorescence intensity per nucleus indicated. **(d)** Quantification of per-nucleus seGFP fluorescence intensity in the absence (−REX3) or presence (+REX3) of REX3, displayed as boxplots. Each dot represents an individual nucleus. Statistical significance was assessed using a two-tailed Student’s t-test (***, p = 0.0001). **(e-f)** Quantification of mean fluorescence intensity in the nucleoplasm of *N. benthamiana* cells expressing the nuclear marker NbH2B-CFP, either uninfected (Control) or invaded by haustoria from transgenic *Phytophthora infestans* lines expressing the secreted effectors SCR74 or Pi22926 fused to mRFP. Nuclei were automatically detected and segmented by HFinder based on the H2B-CFP channel, and the resulting nuclear masks were used to extract fluorescence intensity values from the mRFP channel. Shown are normalized mean intensities for the H2B-CFP **(e)** and effector-mRFP **(f)** channels. Control samples provide background mRFP signal in the absence of effector expression. Dots correspond to individual nuclei, with sample sizes indicated below each condition. Letters denote statistically distinct groups as determined by a Kruskal-Wallis test followed by post hoc comparisons. Ha, haustorium; a.u., arbitrary units.

We then examined whether a comparable quantitative framework could be applied to plant organelles. Using the pTrafficLights reporter system, which co-expresses a secreted GFP (seGFP) and a nuclear-cytoplasmic DsRed from a single *Agrobacterium tumefaciens* T-DNA, we monitored GFP accumulation along the secretory pathway in the presence or absence of the RXLR effector REX3, a known inhibitor of secretion (Evangelisti *et al*., 2017). Nuclei were segmented using HFinder-4 based on DsRed fluorescence **(Fig. 5b-c)**, and the resulting nuclear masks were used to quantify perinuclear GFP signal in the GFP channel. Because all images were acquired under identical imaging conditions, fluorescence intensities could be directly compared across treatments. In agreement with previous observations, REX3 expression resulted in a statistically significant increase in perinuclear GFP signal relative to the control **(Fig. 5d)**, consistent with inhibition of GFP secretion by the effector.

Finally, we asked whether channel-aware segmentation with HFinder could be used to quantitatively capture effector targeting to plant nuclei after secretion from haustoria (Wang *et al*., 2018). We analyzed confocal images of *N. benthamiana* cells expressing the nuclear marker NbH2B-CFP, either alone (Control) or after inoculation with transgenic *P. infestans* strains secreting either the apoplastic protein SCR74 or the cytoplasmic RXLR effector Pi22926 fused to mRFP. Plant nuclei were segmented automatically based on the CFP channel using HFinder-4 **(Fig. 5e, S11**). The resulting nuclear masks were subsequently used to extract fluorescence intensities from the mRFP channel **(Fig. 5f, S11)**. Quantitative analysis revealed a homogeneous accumulation of Pi22926 within the nucleoplasm, whereas SCR74-associated fluorescence remained at background levels comparable to control samples **(Fig. 5f)**. Importantly, these differences emerged from population-level analysis of automatically segmented nuclei, without manual selection of regions of interest. Together, these results demonstrate that channel-aware segmentation with HFinder enables reliable extraction of spatially restricted fluorescence signals from complex infection images, providing a robust and generalizable framework for quantitative analyses at subcellular resolution.

## Discussion

We developed HFinder, a YOLOv8-based framework designed to digitize plant-filamentous microbe interfaces through object-centric detection and delineation of individual infection structures. Trained on a curated dataset of oomycete haustoria labelled via a plant reporter, HFinder robustly detects haustoria across diverse biological contexts, fluorescent reporters and imaging modalities. We demonstrate that the framework generalizes to phylogenetically and morphologically distinct systems, including haustoria from *Phytophthora* species and from downy mildew, and we explore how model tuning can balance specialization against genericity. Beyond haustoria, the same framework can be extended to other discrete biological structures, including hyphal networks and plant organelles. Finally, we illustrate the usefulness of the tool for quantitative analyses at host-pathogen interfaces and in relationships to effector translocation and perturbations of plant cell processes.

Multiple deep learning architectures have been developed for the analysis of biological images. Among them, the U-Net architecture was originally introduced for biomedical image segmentation (Ronneberger *et al*., 2015) and has since been widely adopted across biomedical research (Oskal *et al*., 2019; Huang *et al*., 2019; R *et al*., 2025; Wang *et al*., 2025). The success of U-Net and its derivatives largely stems from pixel-wise segmentation, a strategy where the network predicts a label for every pixel, which has proven particularly effective in biomedical image analysis (Jiangtao *et al*., 2025). Other architectures, such as Mask R-CNN, have also been successfully applied to biology, with tasks as diverse as cell viability assessment and panicle detection (Kong & Chen, 2021; Fan *et al*., 2023; El Akrouchi *et al*., 2025). In contrast to segmentation-first approaches, the YOLO family of algorithms was developed for object detection, with an emphasis on identifying multiple instances within complex scenes while maintaining computational efficiency (Redmon *et al*., 2016). Although initially designed for applications requiring fast inference, YOLO-based approaches have since been successfully applied to fluorescence microscopy, including the detection and tracking of cells and other discrete biological structures in complex backgrounds (Aldughayfiq *et al*., 2023; Stark *et al*., 2023; Krikid *et al*., 2024; Al-Hamadani *et al*., 2025). Here we demonstrate that a lightweight YOLO-based framework can simultaneously detect and delineate intricate host-microbe interfaces and plant organelles in high-magnification confocal images, using limited training data and standard desktop computing resources. While U-Net and related architectures excel at dense pixel-wise segmentation of continuous and homogeneous structures, plant-microbe interfaces are often composed of sparse, heterogeneous and morphologically variable objects. For such configurations, an object-centric detection paradigm is often better suited than pixel-wise segmentation, while offering favorable computational efficiency.

Beyond haustoria, an important perspective of this work lies in its relationship to previous efforts aimed at quantifying plant-microbe interfaces, such as AMFinder (Evangelisti *et al*., 2021). While AMFinder was specifically designed to detect arbuscular mycorrhizal structures using a tile-based segmentation strategy, the object-centric framework implemented in HFinder is, in principle, compatible with the inclusion of arbuscules and other fungal interfaces. This perspective is particularly timely given recent advances in live imaging of arbuscular mycorrhizal fungi, notably through the development of AMSlide, which enables high-resolution, long-term confocal imaging of arbuscule dynamics *in planta* (McGaley *et al*., 2025). In addition, recent work has demonstrated that arbuscules and haustoria can coexist within the same root tissues during dual colonization by mycorrhizal fungi and oomycete pathogens, highlighting the need for image analysis frameworks capable of handling multiple interface types within a single biological context (Guyon *et al*., 2025). More broadly, frameworks such as HFinder may also offer new perspectives for the analysis of fossilized plant-microbe interfaces (Strullu-Derrien *et al*., 2026). Because fossil fungi are primarily defined as morphospecies, quantitative characterization of their interface structures could facilitate more objective comparisons with extant fungal lineages, as well as improved discrimination of fossil taxa.

Extending automated image analysis to complex systems requires approaches that remain compatible with realistic annotation constraints. To reduce manual annotation burden, we explored semi-supervised training strategies based on computer-generated labels derived from masks obtained with intensity-based thresholding. Here, thresholding is deliberately not used to approximate ground-truth annotations, but to provide minimal, low-cost supervision sufficient to bootstrap robust detection models. Similar approaches have been used in microscopy image analysis, where threshold-derived masks serve as a bootstrap for deep learning models and are subsequently improved using learned representations or iterative refinement strategies (Dawoud *et al*., 2023; Xiao *et al*., 2023). This strategy proved sufficient to train robust detection models and reflects a pragmatic compromise between annotation quality and experimental feasibility in realistic experimental settings. Consistent with this rationale, we demonstrate that HFinder can be efficiently adapted to new imaging conditions and biological contexts through transfer learning using limited additional training data, achieving improved detection performance without requiring exhaustive manual annotation. Accordingly, HFinder should be viewed not as a fixed tool optimized for a single interaction, but as a flexible framework that can evolve alongside emerging imaging approaches and increasingly complex biological questions.

Using HFinder, we were able to re-establish through fully automated image analysis several hallmark features of plant-*Phytophthora* interactions that were previously identified using manual inspection. First, we confirmed that the helper NLR NRC4 accumulates preferentially at the haustorial periphery during infection, consistent with earlier observations based on qualitative imaging and manual quantification (Duggan *et al*., 2021). Second, we confirmed the inhibition of GFP secretion mediated by the RXLR effector REX3, a phenotype initially characterized by visual assessment of perinuclear fluorescence patterns and manual intensity measurements (Evangelisti *et al*., 2017). Finally, we showed that the RXLR effector Pi22926, but not the small cysteine-rich protein SCR74, accumulates in plant nuclei following secretion by *Phytophthora infestans*, in agreement with previous reports relying on manual annotation of nuclear signal enrichment (Wang *et al*., 2018). These findings were originally established through careful but low-throughput analyses, often based on representative images or limited numbers of manually quantified regions of interest. By contrast, HFinder enables the same biological conclusions to be derived from large image datasets in a fully automated, unbiased, and statistically scalable manner without manual selection of regions of interest. This shift from manual to object-centric, high-throughput quantification allows infection structures, nuclei, and associated molecular signals to be treated as population-level biological entities rather than illustrative examples. As such, HFinder does not merely reproduce known phenotypes, but demonstrates how automated image analysis can transform established qualitative observations into scalable quantitative readouts, thereby facilitating systematic comparisons across effectors, host genotypes, and infection conditions.

A major remaining challenge in the study of filamentous plant pathogens concerns the biogenesis and dynamics of haustoria. While mature haustoria have been extensively described, their early formation, growth, and morphological transitions *in planta* remain poorly understood, largely due to the difficulty of reliably identifying developing haustoria within complex host tissues (Boevink *et al*., 2020). Addressing these questions will require image analysis frameworks capable of detecting haustoria across developmental stages, including small, transient, or morphologically atypical structures, and operating robustly in live imaging contexts. Although HFinder can already detect branched haustoria in tissues infected with *Phytophthora*, it was not specifically trained to discriminate among multiple haustorial morphotypes. Extending the framework to incorporate morphology-aware classification would enable systematic quantification of haustorial shape distributions within infected tissues. Such capability would be particularly valuable in genetic contexts where haustorial morphology is altered. For example, it would enable the large-scale reassessment of previously reported phenotypes, such as the increased frequency of multilobed haustoria formed by *H. arabidopsidis* in *Arabidopsis* mutants affecting symbiosis-associated SNUPO genes (Ried *et al*., 2019). Automated, morphology-resolved detection would allow these phenotypes to be quantified at scale, their spatial distribution within host tissues to be examined, and haustorial architecture to be related to infection outcomes.

In addition, fast object-centric detection approaches provide a promising foundation for self-driven or adaptive microscopy (Pylvänäinen *et al*., 2023) applied to molecular plant pathology, in which image acquisition could be dynamically guided by automated recognition of nascent infection structures. Beyond two-dimensional analyses, haustoria and associated hyphal networks are inherently three-dimensional structures whose apparent morphology strongly depends on their orientation within host tissues. While volumetric detection in biological imaging has so far been predominantly addressed using U-Net-derived architectures (Çiçek *et al*., 2016), extending object-centric detection to three-dimensional data represents a critical step toward capturing the spatial organization, connectivity, and growth dynamics of invasive hyphae. Together, these perspectives position HFinder as a starting point for integrating quantitative image analysis with live imaging, volumetric microscopy, and ultimately systems-level approaches to host-pathogen interactions.

## Conclusion

In this study, we present HFinder, a deep learning-based framework for the automated detection and segmentation of plant infection structures and organelles in confocal microscopy images. By combining object-centric detection, modest computational requirements and compatibility with low-cost annotation strategies, HFinder enables quantitative, population-scale analyses of plant-microbe interfaces that were previously impractical. This framework thus opens the door to systematic, quantitative comparisons of infection strategies, effector activities and host responses across host genotypes, pathogens and experimental conditions.

## Acknowledgements

We thank Tolga Bozkurt (Imperial College London) for providing the NRC4-GFP construct, Naïma Minet (Sophia Agrobiotech Institute) for maintaining the plants used in this study, and the microscopy platform at the Sophia Agrobiotech Institute for technical assistance. We are grateful to Aurélien Boisson-Dernier (Sophia Agrobiotech Institute) for organizing the stay of Stavros Korovesis. We acknowledge Francine Govers and Tijs Ketelaar (Wageningen University) for insightful discussions.

## Author contributions

S. Korovesis designed the project, acquired and analyzed data, and developed the software. S. Wang, L. Xu and P. Birch generated the effector secretion dataset. I. Giraudon and D. Rosales-Hernandez generated transgenic *Phytophthora* strains. E. Panek and H. Keller provided *Hyaloperonospora* material. L. Boeglin, M.-M. Kostareli, M. Pluis, B. Wang, Y. Wang, and D. Abdennour performed expert annotations. S. Schornack provided the image database used as the training dataset. E. Evangelisti designed the project, acquired and analyzed data, curated the software, secured funding, and wrote the manuscript with contributions from S. Korovesis. All authors read and approved the final manuscript.

