## Supporting Figures 1-11 for "Deep learning enables quantitative subcellular analysis of plant-microbe interfaces"

### Contents

**Figure S1.** Overview of the haustoria training dataset used to develop HFinder-1. 2

**Figure S2.** Training performance of HFinder-1. 3

**Figure S3.** Comparison between HFinder-1 segmentation and expert-derived consensus. 4

**Figure S4.** Training performance of HFinder-2. 5

**Figure S5.** Effect of transfer learning on HFinder performance. 6

**Figure S6.** Transfer learning increases confidence scores for true positive haustorium detections. 7

**Figure S7.** Training dataset used for multiclass HFinder training models. 8

**Figure S8.** Training performance of HFinder-4. 9

**Figure S9.** HFinder-4 frequently merges adjacent chloroplasts into single predictions. 10

**Figure S10.** Weighted scoring and rule-based assignment of predicted classes. 11

**Figure S11.** Nuclear instances used for quantitative analysis of effector nuclear accumulation. 12

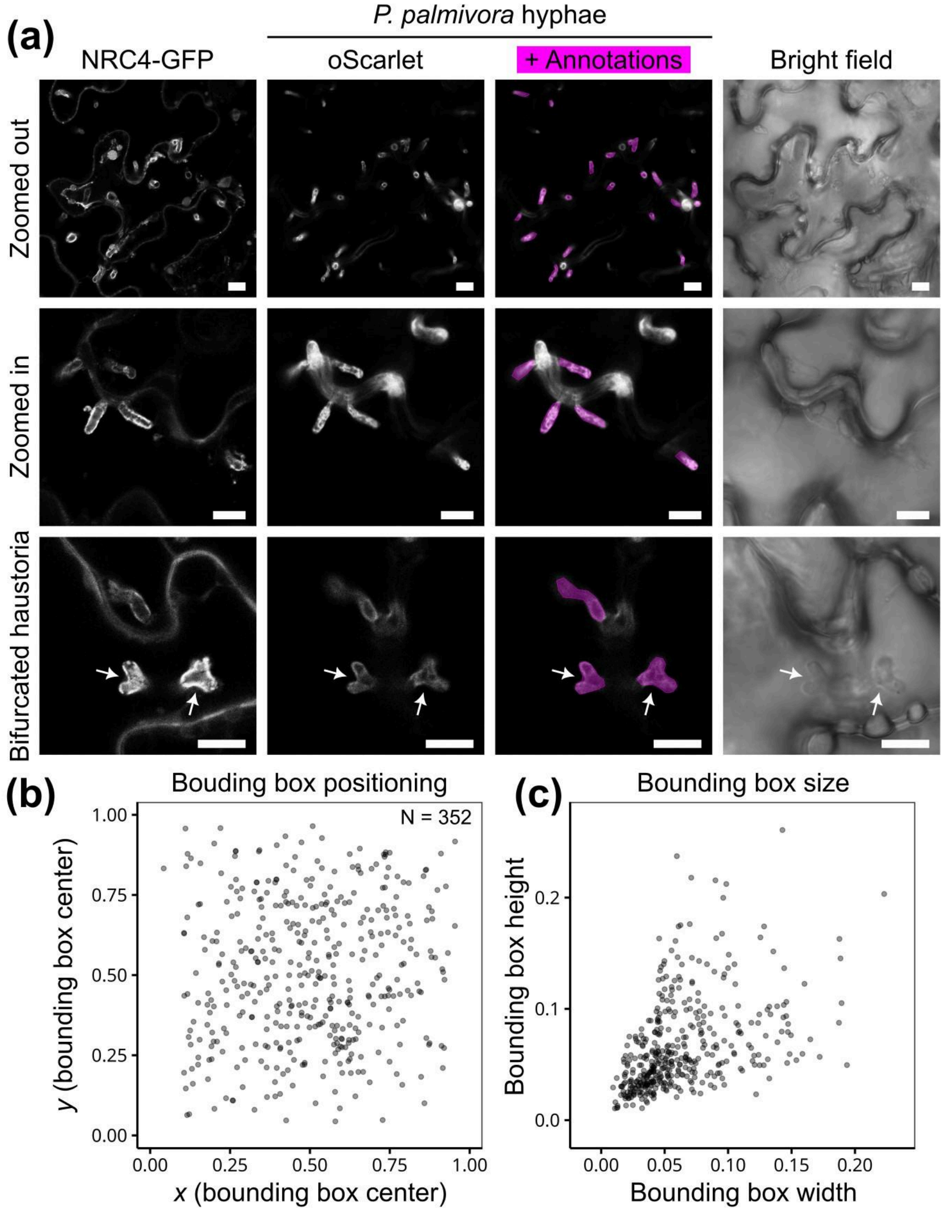


**Figure S1. Overview of the haustoria training dataset used to develop HFinder-1. (A)** Representative confocal images used to assemble the haustoria training set. NRC4-GFP accumulates around haustoria formed by *Phytophthora palmivora* during infection of *Nicotiana benthamiana*, enabling confident identification of bona fide structures. Panels show lower- ("zoomed out") and higher-magnification ("zoomed in") views, as well as examples of bifurcated haustoria (arrows), displayed as raw NRC4-GFP signal, *P. palmivora* hyphae alone or overlaid with manually annotated ground-truth masks (magenta), and the corresponding bright-field images. Scale bars: 10 µm. **(b)** Spatial distribution of annotated haustoria (N = 352) based on the normalized coordinates of bounding-box centers. The homogeneous distribution indicates no positional bias across the field of view. **(c)** Distribution of bounding-box sizes for all annotated haustoria, expressed as normalized width and height. The narrow range of object dimensions reflects the characteristic morphology of *P. palmivora* haustoria within this dataset.

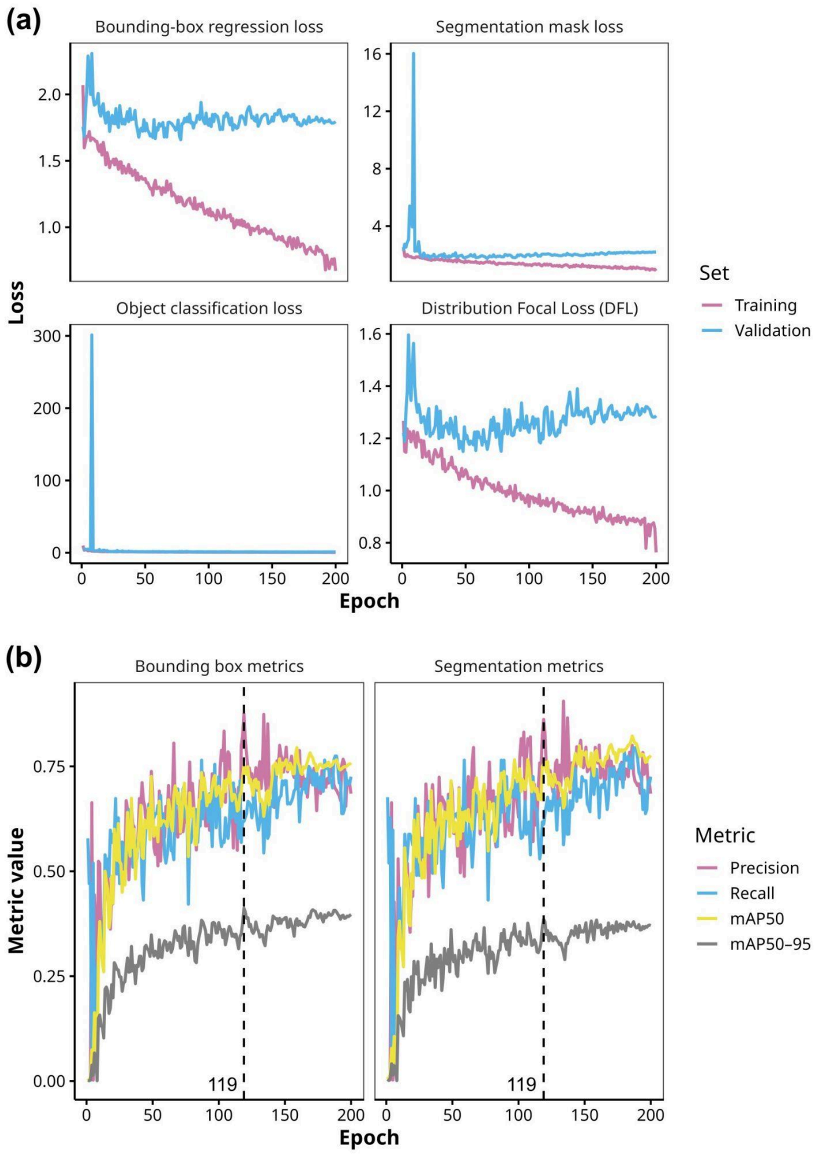


**Figure S2. Training performance of HFinder-1.** HFinder-1 was trained for 200 epochs to detect and segment haustoria using a dataset of 352 manually annotated haustoria. **(a)** Training and validation losses are shown for the four components of the YOLOv8 objective function. Both bounding-box regression and segmentation-mask losses decreased steadily over time, with closely aligned training and validation curves, indicating the absence of overfitting. Object-classification loss rapidly collapsed to zero due to the presence of a single object class. Distribution Focal Loss (DFL), which penalizes inaccuracies in the predicted bounding-box distributions, also showed a consistent downward trend. **(b)** Model generalization was evaluated on the validation set using standard detection metrics: precision, recall, mAP50 (mean average precision at Intersection-over-Union (IoU) = 0.50), and mAP50-95 (mAP averaged over IoU thresholds from 0.50 to 0.95). The optimal checkpoint was reached at epoch 119, corresponding to the maximum YOLOv8 fitness score, defined as a weighted combination of mAP50 and mAP50-95.

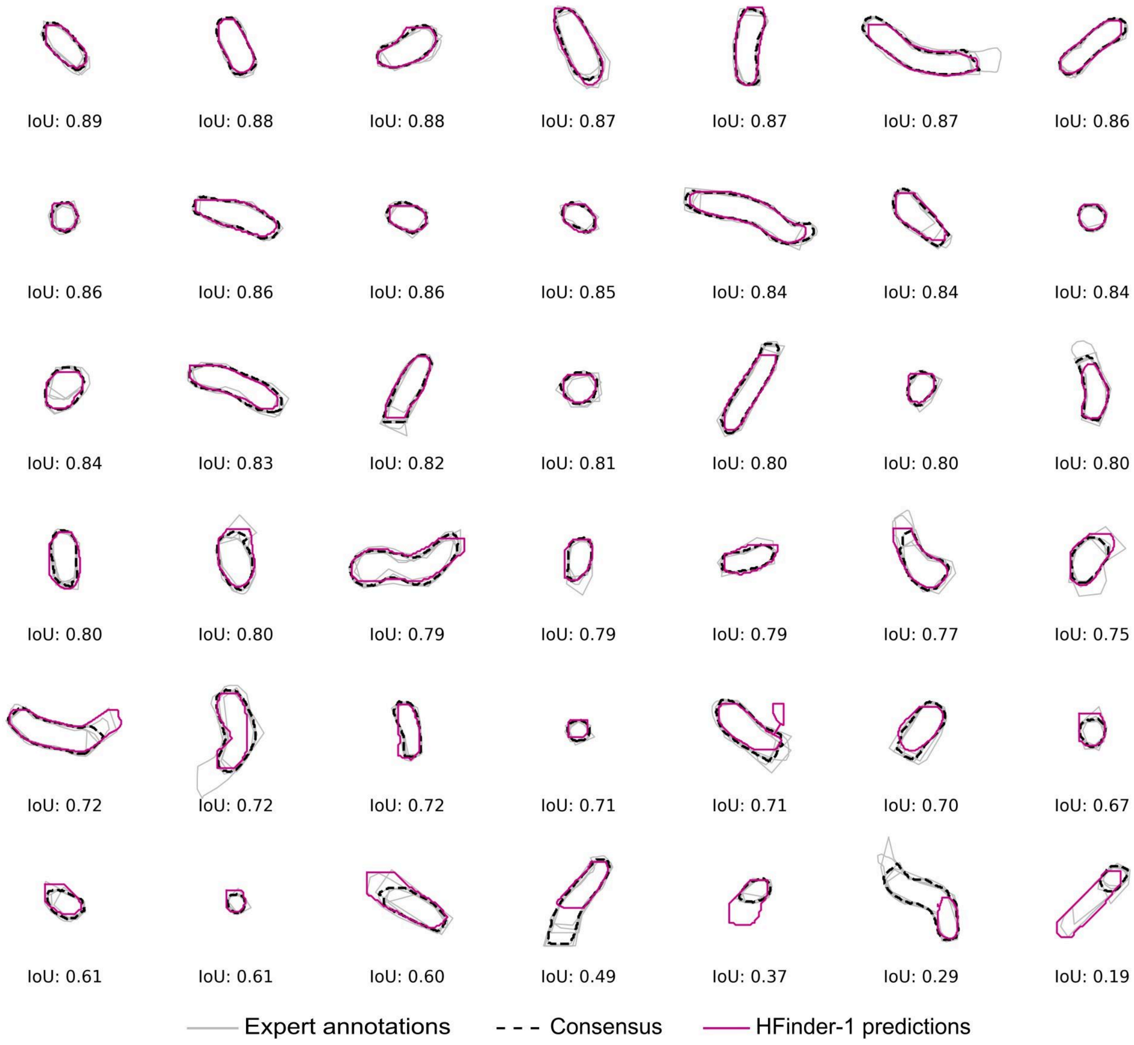


**Figure S3. Comparison between HFinder-1 segmentation and expert-derived consensus.** A set of 48 NRC4-labelled haustoria, sampled across different magnifications and spatial orientations, were manually segmented by six independent annotators using the Makesense.ai platform (light-gray polygons). These annotations were integrated into a consensus mask (black dashed polygons), which was compared to HFinder predictions (magenta outlines). Each panel shows one haustorium, and the panels are sorted in decreasing order of intersection-over-union (IoU) between the consensus and the model prediction. Five haustoria that the model did not detect (false negatives) are not shown. Haustoria are not displayed to scale.

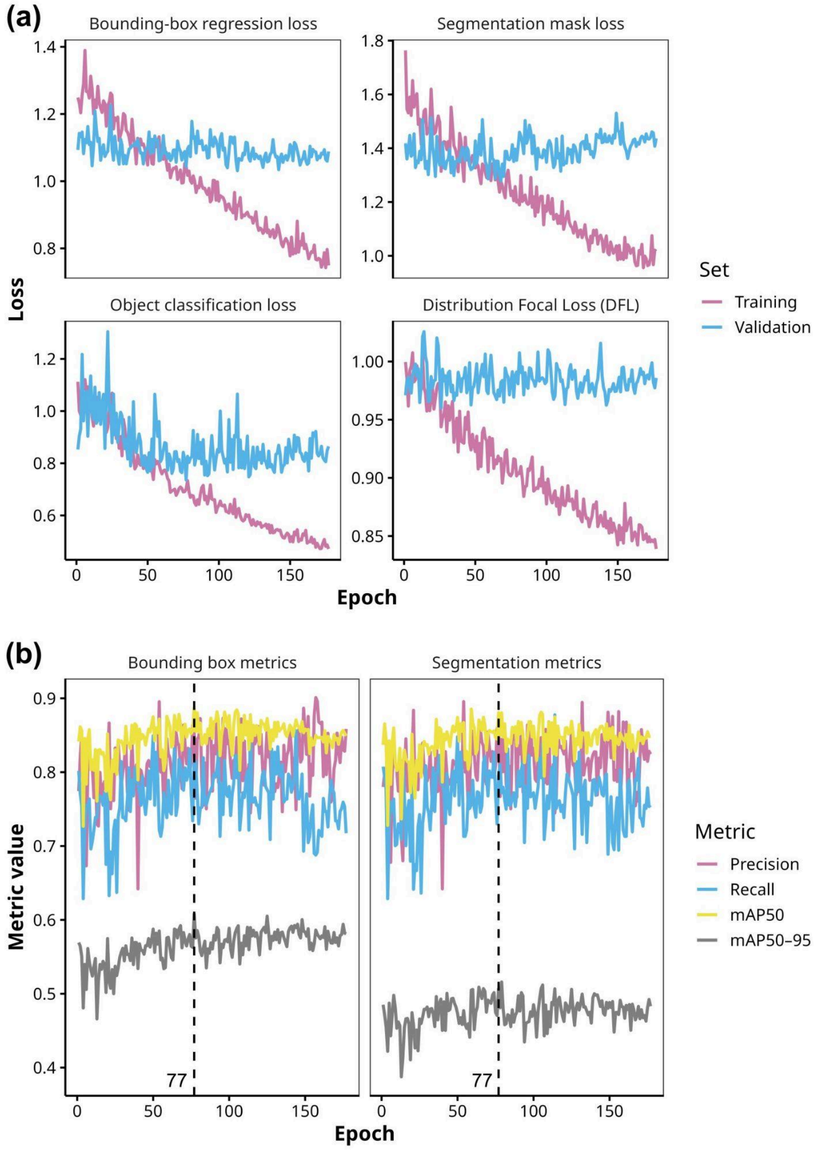


**Figure S4. Training performance of HFinder-2.** HFinder-2 was trained for 200 epochs using an expanded dataset combining the 352 manually annotated haustoria used for HFinder-1 with 496 additional haustoria curated through supervised annotation with HFinder-1. Training was initialized from the best HFinder-1 weights to promote cross-condition generalization. **(a)** Loss curves for the four components of the YOLOv8 objective function illustrate stable convergence throughout training, consistent with effective fine-tuning on the augmented dataset. **(b)** Validation metrics indicate improved performance stability relative to HFinder-1, with the highest YOLOv8 fitness score achieved at epoch 77, which was selected as the optimal checkpoint for downstream analyses.

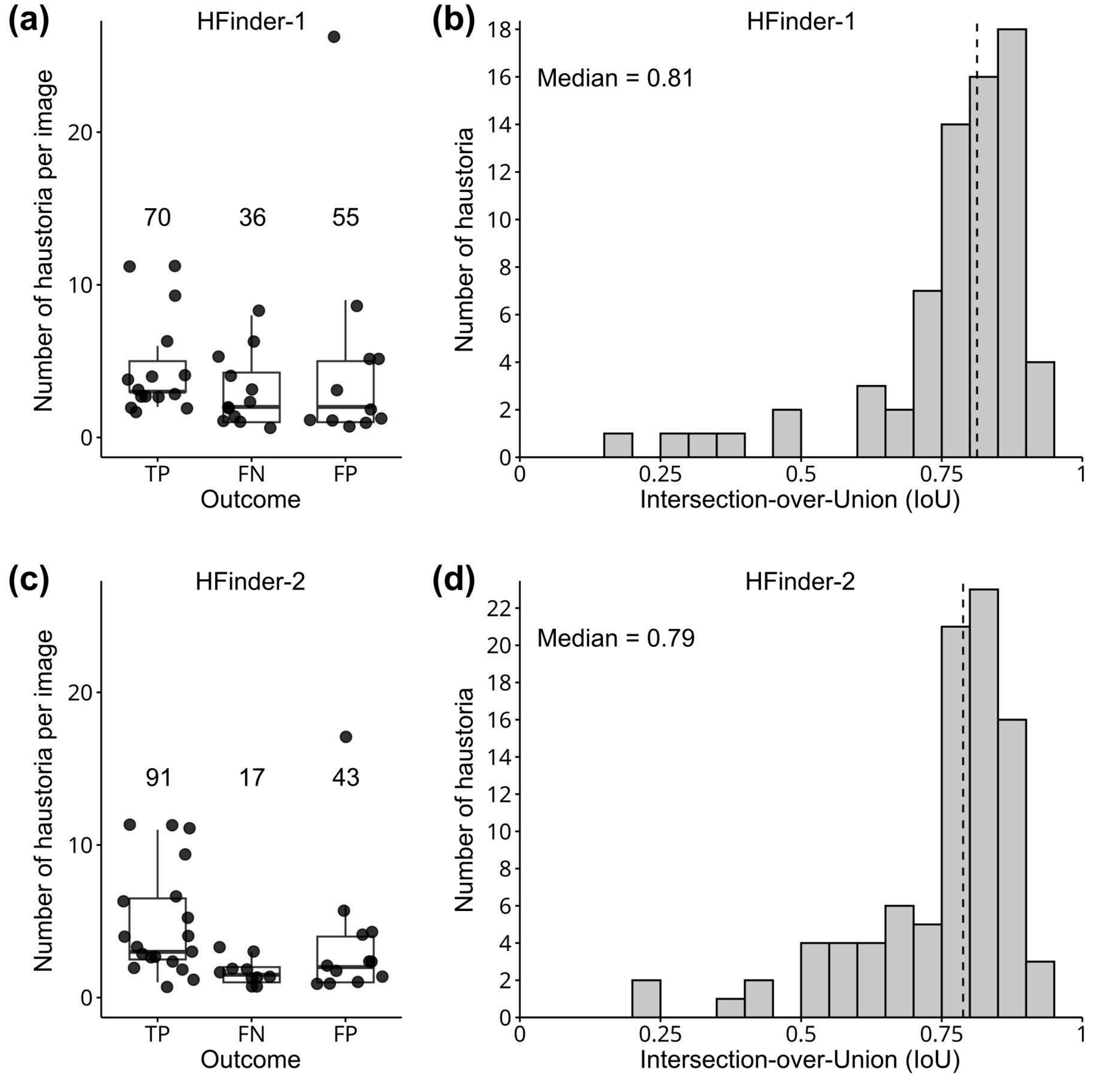


**Figure S5. Effect of transfer learning on HFinder performance. (a-d)** HFinder was used to predict haustoria on a test set of 18 images, comprising the 9 images previously used for validating HFinder-1 and 9 images representative of the dataset used for transfer learning. Boxplots showing the number of true positives (TP), false negatives (FN), and false positives (FP) per image are shown for HFinder-1 **(a)** and HFinder-2 **(c)**. The distribution of intersection-over-union (IoU) values between predicted haustoria segmentations and expert consensus annotations is shown for HFinder-1 **(b)** and HFinder-2 **(d)**.

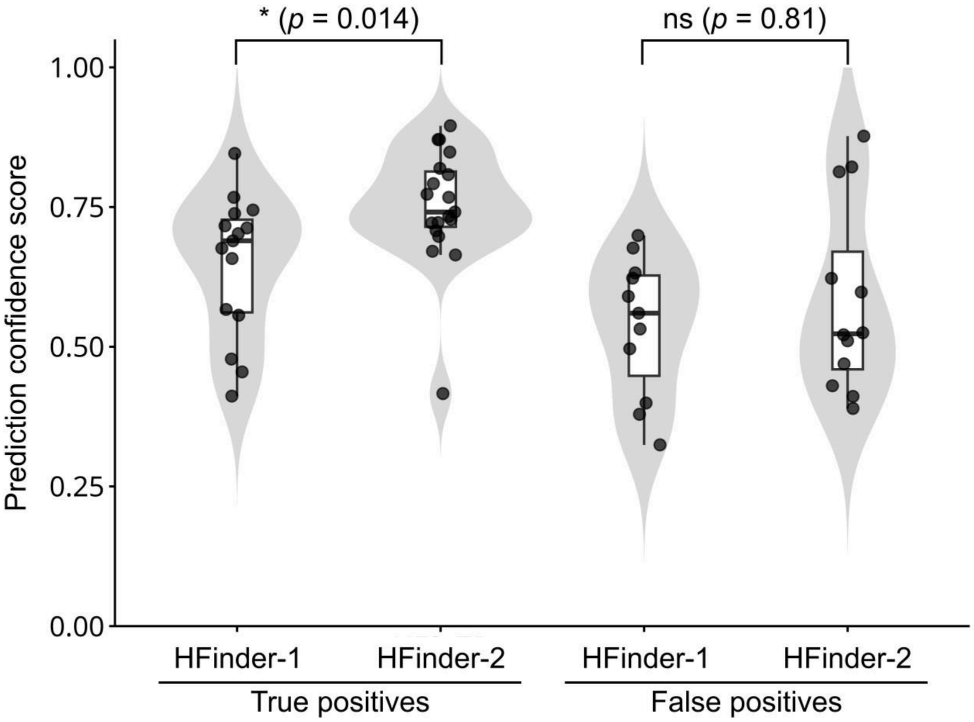


**Figure S6. Transfer learning increases confidence scores for true positive haustorium detections.** Violin plots showing the distribution of YOLO prediction confidence scores for true positive (left) and false positive (right) haustorium detections obtained with HFinder-1 and HFinder-2. Each point represents the confidence score of a single detected instance; boxplots indicate the median and interquartile range. Statistical significance was assessed using a paired Wilcoxon signed-rank test. ns: not significant.

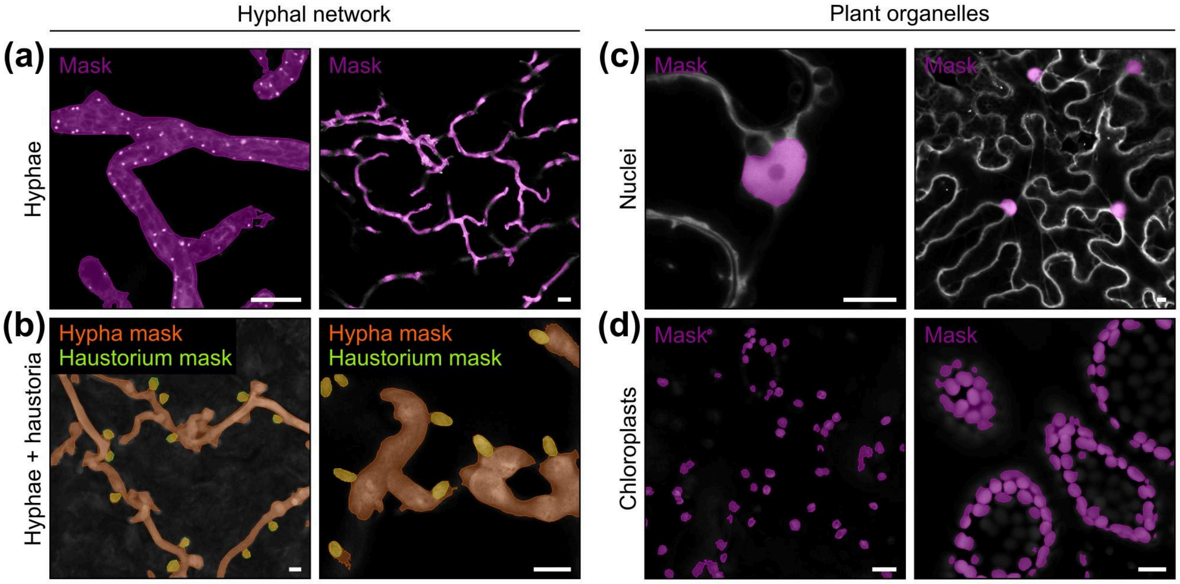


**Figure S7. Training dataset used for multiclass HFinder models. (a-d)** Representative confocal images illustrating the diversity of imaging conditions and object classes used to train HFinder-4. Images include hyphal networks grown under axenic conditions **(a)**, hyphae bearing haustoria during plant infection **(b)**, and plant organelles, including nuclei **(c)** and chloroplasts **(d)**. Segmentation masks are overlaid in magenta for single-class predictions and in orange and yellow for co-occurring classes within the same image. Scale bars: 10 µm.

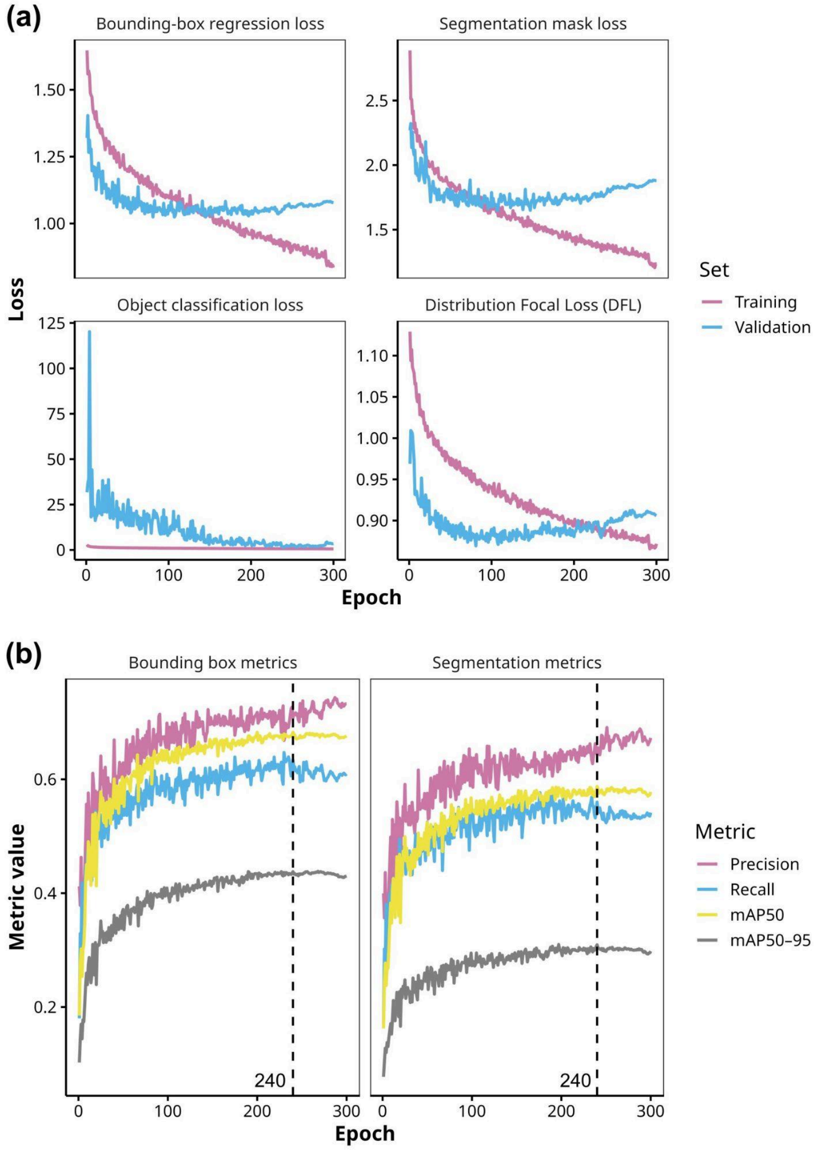


**Figure S8. Training performance of HFinder-4.** HFinder-4 was trained for 300 epochs using a dataset comprising manually annotated images featuring hyphae, haustoria, nuclei, and chloroplasts. Training was initialized from the naive YOLOv8 weights. **(a)** Loss curves for the four components of the YOLOv8 objective function illustrate stable convergence throughout training, consistent with effective fine-tuning on the augmented dataset. **(b)** Validation metrics indicate improved performance stability relative to HFinder-4, with the highest YOLOv8 fitness score achieved at epoch 240, which was selected as the optimal checkpoint for downstream analyses.

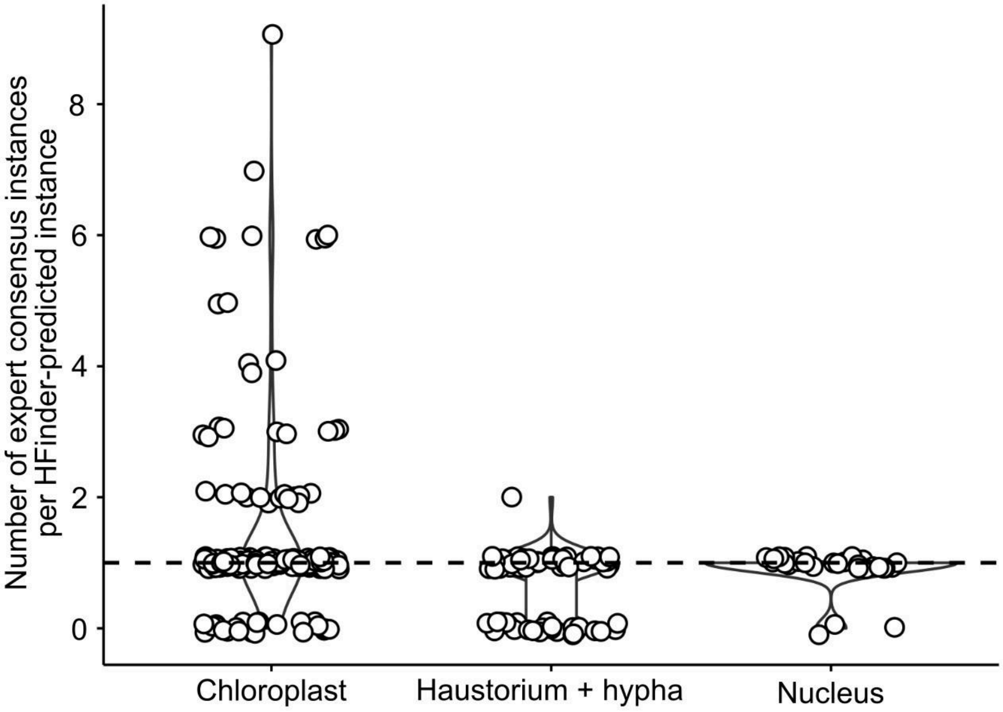


**Figure S9. HFinder-4 frequently merges adjacent chloroplasts into single predictions.** Violin plots showing the number of expert consensus instances associated with each HFinder-4-predicted instance, for chloroplasts (left), hyphae with haustoria (middle), and nuclei (right). Values greater than 1 indicate merging of multiple expert-annotated instances into a single HFinder prediction. Values equal to 1 correspond to one-to-one matches, whereas values of 0 indicate false-positive predictions or strong mismatches between expert annotations and model outputs.

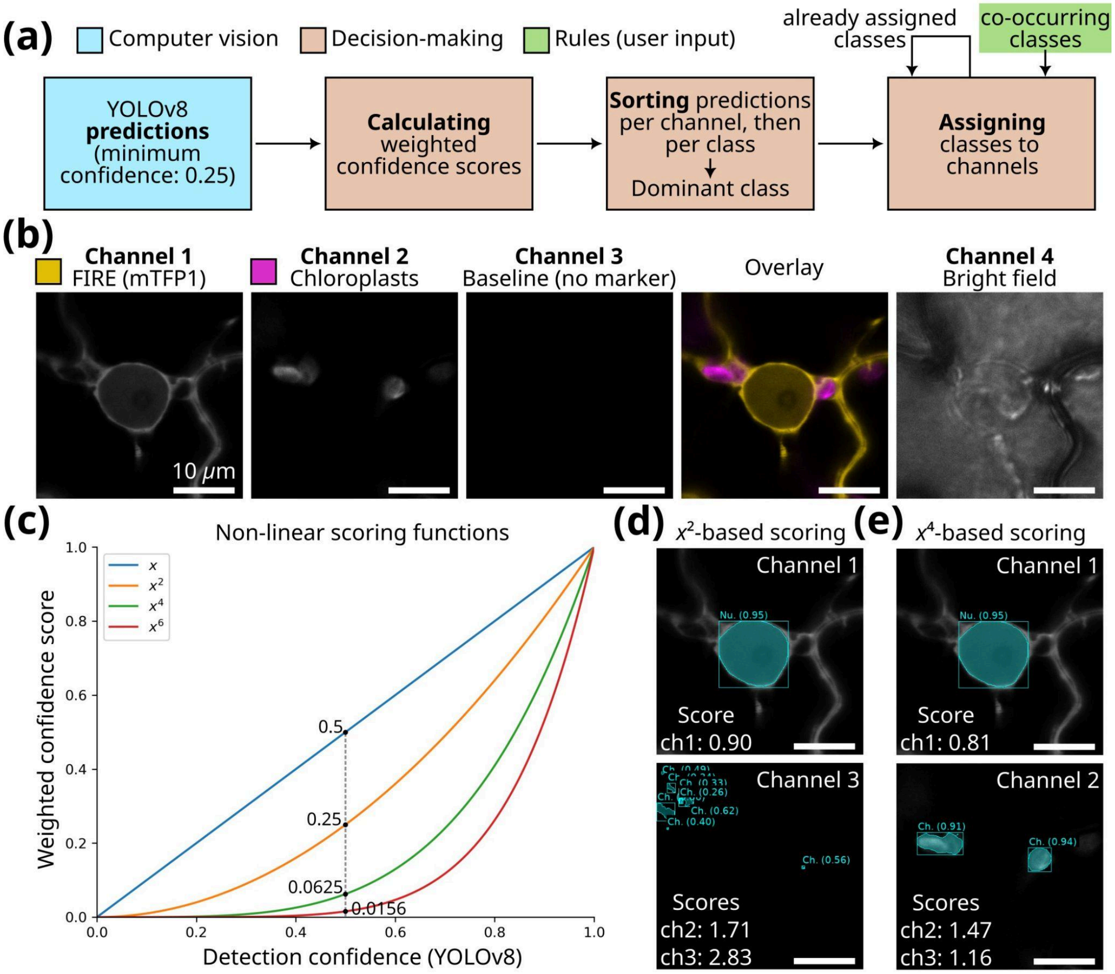


**Figure S10. Weighted scoring and rule-based assignment of predicted classes. (a)** Schematic overview of the decision-making pipeline. YOLOv8 generates object predictions from each microscopy channel (computer vision). Confidence values are aggregated into weighted class scores using non-linear functions and used for sorting by channel and by class, enabling automatic assignment of the dominant class to a channel, with optional co-dominant classes included based on user-defined rules (Rules, green). Classes that are already assigned are excluded from further use (Decision-making). **(b)** Example raw data illustrating four imaging channels: mTFP1-labeled FIRE effector, chloroplast autofluorescence, a baseline channel with no specific marker, and brightfield. **(c)** Weighted scoring functions used to calculate class scores. Non-linear transformations of detection confidence ($x$, $x^2$, $x^4$, $x^6$) reduce the influence of weak predictions while emphasizing strong ones. For example, a detection confidence of 0.5 contributes 0.5, 0.25, 0.0625, or 0.0156 to the class score depending on the chosen exponent. **(d)** Example with quadratic scoring. Weak but numerous detections in the baseline channel (channel 3) yield a higher total score than the signal in channel 2, leading to a misassignment. **(e)** Example with quartic scoring. High-quality detections dominate the score, suppressing background noise and correctly identifying the dominant classes in channels 1 and 2. Scale bars: 10 μm.

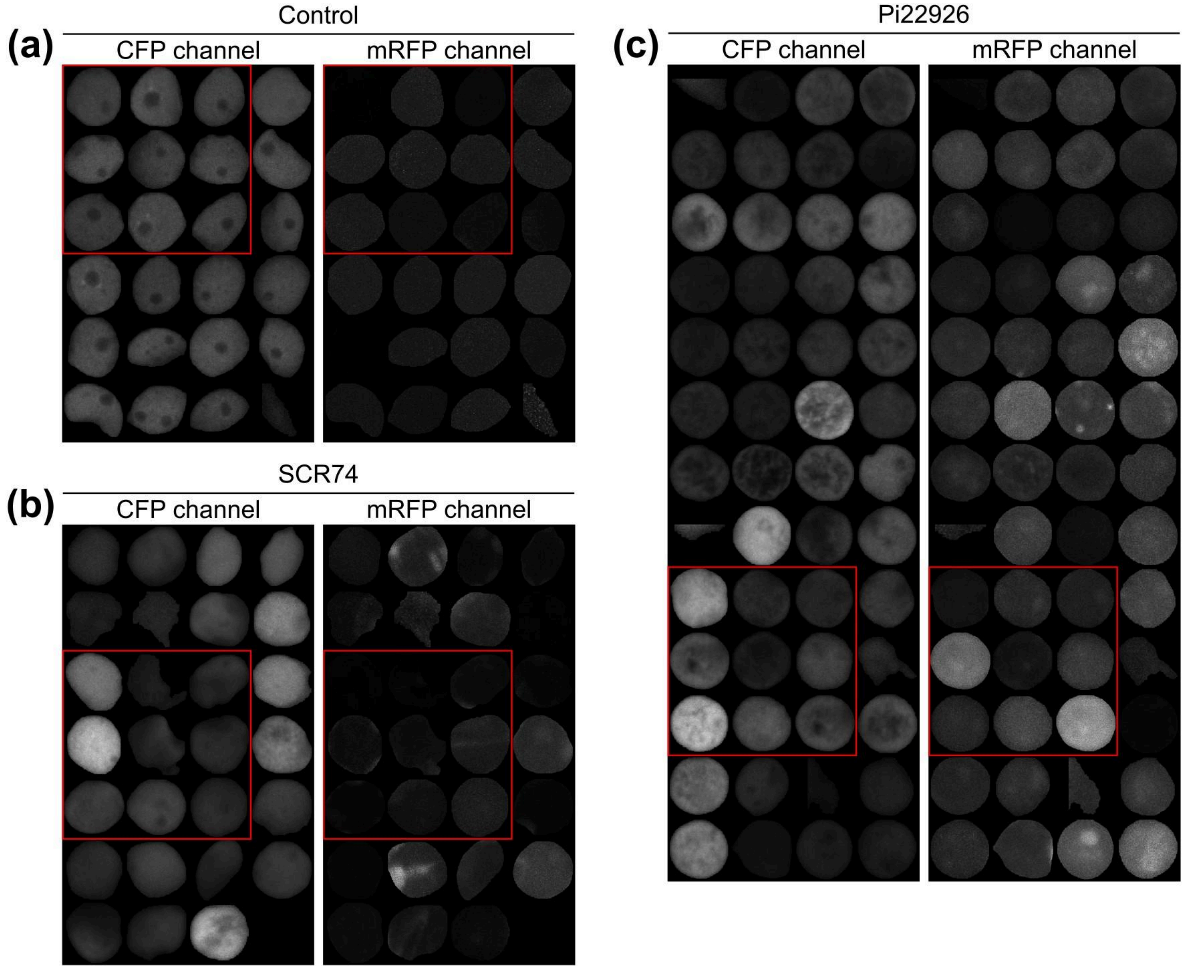


**Figure S11. Nuclear instances used for quantitative analysis of effector nuclear accumulation.** Representative montages showing all individual plant nuclei included in the quantification of effector accumulation in nuclei after secretion from *Phytophthora*. Nuclei were segmented based on the NbH2B-CFP channel (left panels), and corresponding fluorescence intensities were extracted from the mRFP channel (right panels). Panels show nuclei from control samples lacking mRFP-labelled effectors, which therefore report background fluorescence levels **(a)**, samples infected with *P. infestans* secreting the apoplastic protein SCR74 **(b)**, and samples infected with *P. infestans* secreting the cytoplasmic RXLR effector Pi22926 **(c)**. Each tile corresponds to a single segmented nucleus. Images are displayed with identical intensity scaling within each channel. Red frames indicate the nuclei shown in Figure 5.
